# Opposing transcription factors MYCL and HEY1 mediate the Notch-dependent airway stem cell fate decision

**DOI:** 10.1101/2022.10.05.511009

**Authors:** Lauren E. Byrnes, Rachel Deleon, Jeremy F. Reiter, Semil P. Choksi

**Affiliations:** Department of Biochemistry and Biophysics, Cardiovascular Research Institute, University of California, San Francisco, CA 94158, USA; Chan Zuckerberg Biohub, San Francisco, CA 94158, USA

**Keywords:** Basal stem cell, multiciliated cell, secretory cell, differentiation, cell fate, transcriptional regulation

## Abstract

Tissue function depends on the relative proportions of multiple cell types. In the airway, basal stem cells differentiate into both multiciliated and secretory cells, which together protect the lungs from inhaled pathogens and particulates. To define how airway stem cells differentiate, we mapped differentiation trajectories using single-cell mRNA sequencing (scRNA-seq) and identified a transitional intermediate cell state in between basal stem cells and differentiated cells. These intermediate cells induce different gene expression programs that precede differentiation into either multiciliated or secretory cells. For example, we found that within the intermediate cell population, multiciliated cell precursors express *Mycl*, encoding a MYC-family transcription factor, and secretory cell precursors express *Hey1*, encoding a transcriptional repressor. We also found that Notch signaling acts on intermediate cells to repress *Mycl* and induce *Hey1*. We further show MYCL expression is sufficient to drive multiciliated cell fate, whereas HEY1 expression is sufficient to repress multiciliated cell fate. Using CUT&RUN, we made the surprising observation that MYCL and HEY1 bind to many of the same regulatory elements near genes encoding early regulators of multiciliated cell differentiation. We conclude that intermediate cells receiving Notch signals induce HEY1 to repress the multiciliated cell fate and become secretory cells, while intermediate cells not receiving Notch signals induce MYCL to promote the multiciliated cell fate. These experiments reveal that during airway stem cell differentiation Notch signaling balances the production of two different cell types by regulating the functions of two opposing transcription factors, MYCL and HEY1.

## Introduction

The conducting airways bring air to the distal lung and generate a mucociliary barrier that captures and removes inhaled particulates and respiratory pathogens. The luminal layer of the airway epithelium consists of secretory cells which produce mucus and multiciliated cells which sweep mucus away from the lower airway. Below these luminal cells lies a layer of basal stem cells which both self-renew and give rise to mature multiciliated and secretory cells (Hackett et al., 2008; Hong et al., 2004; Rock et al., 2009).

How basal stem cells give rise to two differentiated cell types in defined proportions remains unclear. The ratio of multiciliated to secretory cells is critical for airway function, exemplified by the increased proportion of secretory cells in chronic respiratory diseases such as asthma and chronic obstructive pulmonary disease (COPD). This skewed ratio leads to excess mucus production and reduced mucus clearance, which contribute to mucus plugs that can be fatal in severe respiratory disease (Erle and Sheppard, 2014; Tilley et al., 2015). Isolated disruption of multiciliated cell function causes a different human disease called primary ciliary dyskinesia (PCD), characterized by recurrent respiratory infections and bronchiectasis (Horani and Ferkol, 2021; Legendre et al., 2021).

The key regulator of cell fate choice in the airway is Notch signaling: Notch activation promotes the formation of secretory cells, and Notch inhibition promotes the formation of multiciliated cells (Guseh et al., 2009; Morimoto et al., 2010; Tsao et al., 2009). Overactivation of Notch signaling is associated with the secretory cell hyperplasia observed in respiratory disease (Danahay et al., 2015; Kang et al., 2009; Pardo-Saganta et al., 2015), suggesting that attenuating Notch signaling could be an important therapeutic approach to correcting cell fate proportions in the diseased airway (Lafkas et al., 2015). Therefore, we have sought to understand how Notch signaling directs secretory and multiciliated cell fates.

Downstream of Notch signaling, multiciliated cell fate depends on two Geminin-family transcriptional regulators, GMNC (Geminin Coiled-Coil Domain Containing) and MCIDAS (Multiciliate Differentiation And DNA Synthesis Associated Cell Cycle Protein), both induced early during multiciliated cell differentiation (Arbi et al., 2016; Boon et al., 2014; Lu et al., 2019; Ma et al., 2014; Stubbs et al., 2012; Terré et al., 2016). GMNC and MCIDAS induce the expression of downstream transcription factors such as MYB and FOXJ1 (Forkhead Box J1) as well as other proteins such as CCNO (Cyclin O) that promote multiciliated cell maturation (Brody et al., 2000; Chen et al., 1998; Choksi et al., 2014b, 2014a; Funk et al., 2015; Pan et al., 2014; Tan et al., 2013).

Multiciliated cell differentiation requires the synthesis of hundreds of basal bodies, each of which extends a motile cilium at the apical surface of the cell. The complexity of multiciliated cell lineage differentiation has been broken down into four main stages based on the expression and localization of proteins required to make multiple cilia (Tan et al., 2013; Vladar and Stearns, 2007). During Stage I, basal body biogenesis proteins are synthesized. During Stage II, basal bodies are generated. During Stage III, basal bodies migrate and dock to the apical membrane. Finally, at Stage IV, basal bodies generate motile cilia. During Stages I-III, these cells possess deuterosomes, transient structures involved in basal body generation (Sorokin, 1968). How Notch signaling controls the transition of basal stem cells to MCIDAS- and MYB-expressing deuterosomal cells for the eventual formation of mature multiciliated cells is not understood.

Profiling transcriptomes at single-cell resolution allows for detailed charting of the expression changes corresponding to cellular differentiation (Soldatov et al., 2019). We have used single-cell mRNA sequencing (scRNA-seq) to map airway basal stem cell differentiation. The resultant developmental trajectories identified an intermediate cell state characterized by erasure of the stem cell transcriptional program but preceding the induction of genes characteristic of early multiciliated and secretory cell differentiation. We generated CRISPR knockouts of *Gmnc* and *Mcidas* and used scRNA-seq to identify how these transcriptional regulators direct the transition from intermediate cells to multiciliated cell precursors. Within different subpopulations of intermediate cells, we identified expression of two transcription factors, MYCL and HEY1. We identified MYCL as an early driver of multiciliated cell differentiation, whereas HEY1 is an early repressor of the multiciliated cell fate expressed in secretory cell precursors. Within the intermediate cell population, Notch signaling activates HEY1 and represses MYCL. Together, these experiments reveal how Notch acts on differentiating basal stem cells to regulate early transcriptional effectors and drive alternative cell fates, critical for defining the cell composition of the airway.

## Results

### Basal stem cells adopt an intermediate cell state before differentiating along a defined lineage

Primary mouse tracheal epithelial cells (mTECs) can generate a pseudostratified epithelium similar to that of the upper airway, containing basal stem cells and luminal differentiated cells (You and Brody, 2013). To define transitional states during basal stem cell differentiation, we assessed transcriptomes of differentiating mTECs using scRNA-seq. Specifically, we generated three independent scRNA-seq datasets of mTECs differentiated for three days at air/liquid interface. To ensure capture of all stages of multiciliated cell differentiation, we also generated a dataset enriched for multiciliated lineage cells by fluorescence activated cell sorting (FACS) of red fluorescent mTECs cultured from *Foxj1-CreERT Rosa-tdTomato* transgenic mice (Figure 1A and S1A, see materials and methods). Integrating scRNA-seq data from all datasets captured 21,740 cells and identified 8 UMAP clusters (Figure 1A and S1B). This dataset will be available for query on the CellxGene scRNA-seq platform (https://cellxgene.cziscience.com/collections/).

**Figure 1.**
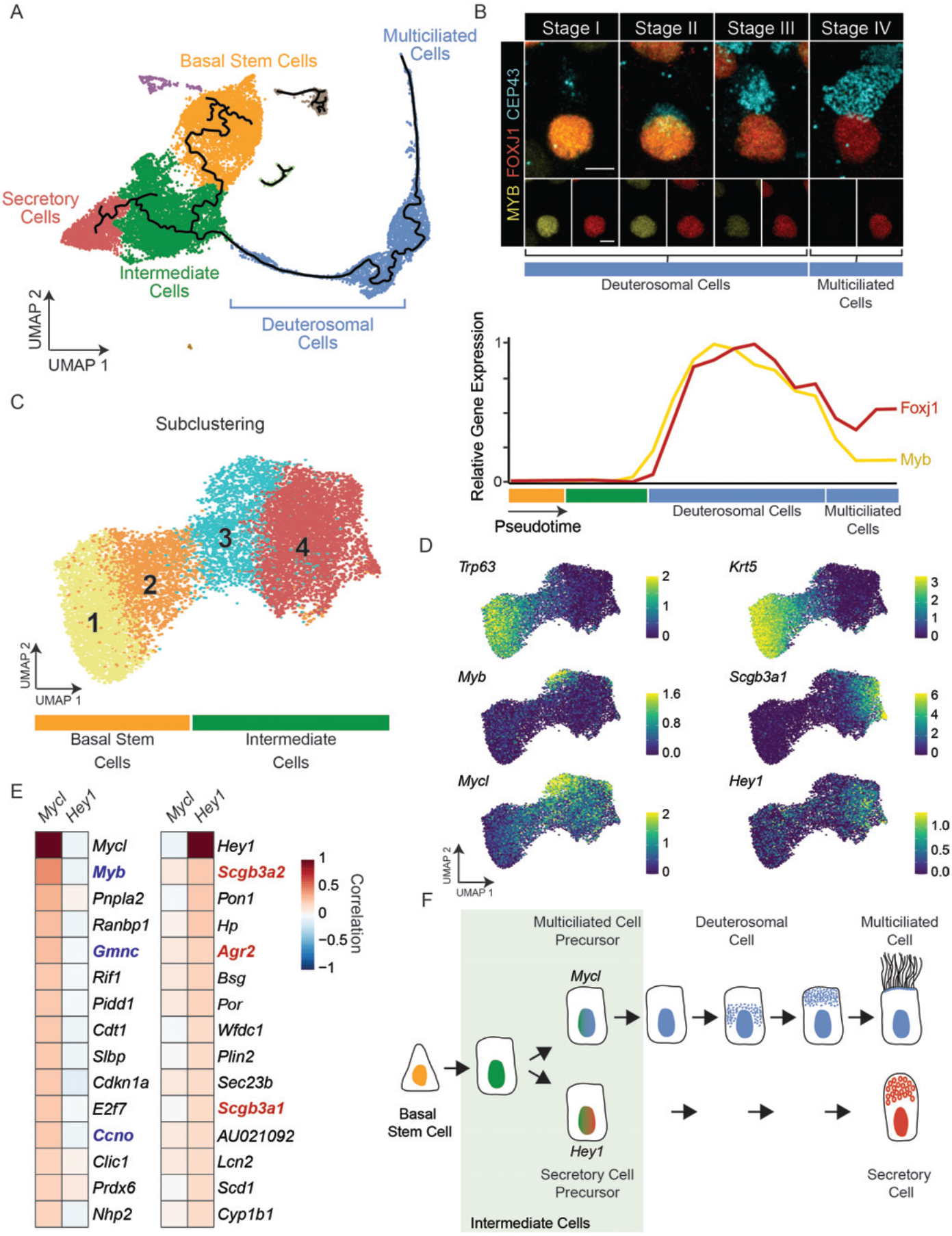
Identification of an intermediate cell state bridging basal stem cell differentiation into secretory and multiciliated lineages. A) UMAP of scRNA-seq data generated from primary mouse tracheal epithelial cells three days after differentiation. Cells are colored by cluster identity. Pseudotime paths are denoted as black lines. B) Stages of multiciliogenesis visualized by immunofluorescence for basal bodies (CEP43), MYB and FOXJ1 (top panel). Scale bars represent 5 μm. Normalized expression of *Myb* and *Foxj1* over the pseudotime path from basal stem cells to multiciliated cells depicted in (A) (bottom panel). C) UMAP of subclustered basal stem cells and intermediate cells from primary mouse tracheal cell scRNA-seq from (A). Cells are colored by cluster identity. D) Expression of select marker genes overlaid on the UMAP of subclustered basal stem cells and intermediate cells. Heat map of the top 15 correlated genes with either *Mycl* or *Hey1* within the basal and intermediate cells. Known regulators of multiciliated cell differentiation are highlighted in blue and known regulators of secretory cell differentiation are highlighted in red. F) Model of basal stem cell differentiation through an intermediate cell state to a multiciliated or secretory cell fate. A subset of intermediate cells expresses *Mycl* along with other multiciliated cell genes or Hey1 with other secretory cell genes.

Comparison to previously described cell type-specific markers identified populations of basal stem cells (e.g., expressing *Trp63*) and secretory cells (e.g., expressing *Scgb3a1*) (Figure S1C). Multiciliated cell differentiation, defined by expression of multiciliated lineage transcription factors *Myb* and *Foxj1* (Brody et al., 2000; Chen et al., 1998; Pan et al., 2014; Tan et al., 2013), was represented in the UMAP by a long arcing cluster (Figure 1A and S1C). Other clusters represented less prevalent cell types, such as neuroendocrine or proliferating cells (Figure S1D and Supplementary Table 1).

Pseudotime analysis predicted a differentiation trajectory starting at basal stem cells and splitting within an intermediate cell population, with one fork proceeding through the multiciliated lineage cluster (Figure 1A). We hypothesized that the long arc of the multiciliated lineage cluster represents a continuum of related transcriptional states describing multiciliated cell differentiation. MYB and FOXJ1 are expressed throughout multiciliated cell differentiation (Figure 1B). Both MYB and FOXJ1 expression are observed during the deuterosomal cell phase (Stages I-III) of the multiciliated lineage while FOXJ1 is maintained in mature multiciliated cells (Stage IV, Figure 1B). Consistent with the expression dynamics of the proteins, the scRNA-seq data suggested that *Myb* is expressed early in the multiciliated cell trajectory and downregulated in mature multiciliated cells while *Foxj1* expression is maintained in mature multiciliated cells (Figure 1B).

In further support of the hypothesis that the arc of the multiciliated cell trajectory represents differentiation, *Deup1*, encoding the principal deuterosome component (Zhao et al., 2013), was expressed early in the multiciliated cell trajectory and genes encoding dynein components required for ciliary motility (e.g., *Dnah5*) were expressed late (Figure S1D). Thus, pseudotime analysis and analysis of expression of previously characterized genes indicate that the multiciliated cluster reflects stages of differentiation, from deuterosomal stages through ciliogenesis.

An additional large cell cluster was positioned between the basal stem cells and the multiciliated and secretory cell clusters in the UMAP (Figure 1A). This intermediate cluster was not enriched for expression of previously identified markers. To determine whether this intermediate cluster represents a previously defined cell type, we used label transfer to compare our scRNA-seq data to previously generated human airway scRNA-seq data (Deprez et al., 2020). Comparing these datasets confirmed the identity of basal stem, secretory and multiciliated cell clusters but did not reveal a previously defined descriptor for the intermediate cluster (Figure S1E).

Plotting the expression of markers of basal stem cells (*Trp63*), multiciliated cells (*Myb*, *Foxj1*), and secretory cells (*Scgb3a1*) highlighted the placement of the intermediate cluster at the nadir of the intersection of gradients of basal stem, multiciliated, and secretory cell transcriptional identities (Figure S1C). Based on the UMAP position of the intermediate cluster between the basal stem cells and the differentiating lineages, we hypothesized that this cluster represents a transitory intermediate state. In support of this this hypothesis, the basal stem cell pseudotime trajectory branched within this intermediate cluster to lead to the multiciliated or secretory cell clusters (Figure 1A). These data support the idea that the intermediate cells, lacking a distinct transcriptional signature (Figure S1F), are in a transient state that can adopt either a secretory or multiciliated cell fate.

### Intermediate cells initiate transcriptional programs that presage different lineages

To better characterize the relationship of basal stem cells to this intermediate population, we subclustered basal stem and intermediate cells together, revealing four subclusters (Figure 1C). Assessing the expression of basal stem cell, multiciliated cell, and secretory cell markers in the four subclusters revealed that subclusters 1 and 2 express high levels of *Trp63* and *Krt5* and therefore represent the basal stem cells (Figure 1D). Subclusters 3 and 4 lack expression of both *Trp63* and *Krt5*, and therefore represent intermediate cells. To assess whether intermediate cells initiate differentiation programs, we assayed for the expression of multiciliated and secretory cell genes in subclusters 3 and Subcluster 3 exhibited low expression of *Myb*, a marker of the early multiciliated cell lineage, and subcluster 4 exhibited low expression of *Scgb3a1*, a marker of the secretory cell lineage (Figure 1D).

To identify other genes induced in intermediate cells, we conducted differential gene expression analysis between subclusters 3 and 4. *Mycl* (*MYCL proto-oncogene, bHLH transcription factor*) expression was enriched in subcluster 3 (Supplementary Table 2). *Mycl* expression was broader than, but overlapped with, *Myb* expression (Figure 1D). To better define, independent of clustering, genes co-expressed with *Mycl*, we used gene correlation analysis. We identified 49 genes uniquely correlated with *Mycl* (R^2^ > 0.15) (Figure 1E, Supplementary Table 3), which included multiple genes involved in early multiciliated cell differentiation such as *Gmnc*, *Myb* and *Ccno*. These data suggest *Mycl* is expressed in multiciliated cell precursors within the intermediate population.

In addition to *Scgb3a1*, Subcluster 4 was enriched for other secretory markers, including *Scgb1a1* and *Muc5b* (Supplementary Table 2), suggesting it represents secretory cell precursors within the intermediate population. These expression analyses raise the possibility that multiciliated and secretory cell lineages emerge from the intermediate cell state.

Notch signaling promotes the production of secretory cells and inhibits the production of multiciliated cells (Rock et al., 2011; Tsao et al., 2009). To begin to determine if Notch signaling acts within basal stem cells or intermediate cells to discriminate between these fates, we examined the expression of Notch pathway genes in basal stem cells and intermediate cells. We found that *Notch1* and *Notch2*, encoding receptors, are expressed in basal stem cells and intermediate cells, respectively (Figure S2A). Notably, expression of *Hey1*, a Notch target gene encoding a bHLH transcriptional repressor (Dunwoodie et al., 2002), is enriched in intermediate cell subcluster 4 together with *Scgb3a1* (Figure 1D). Gene correlation identified 14 genes that uniquely correlated with *Hey1* (R^2^ > 0.15) (Figure 1E), which include multiple genes involved in secretory cell function (e.g., *Agr2*, Chen et al., 2009). Thus, *Hey1* may be a Notch target induced within the intermediate cell population in secretory cell precursors. These data raise the possibility that *Mycl*-expressing intermediate cells are multiciliated cell precursors and *Hey1*-expressing intermediate cells are secretory cell precursors (Figure 1F).

If *Mycl* expression defines multiciliated cell precursors and *Hey1* expression defines secretory cell precursors, then the expression of these genes should be anti-correlated. Indeed, *Mycl* and *Hey1* expression was anti-correlated in scRNA-seq data (Figure S2B). Multiplexed RNA fluorescent *in situ* hybridization (RNA-FISH) confirmed the segregation of *Mycl* and *Hey1* transcripts in distinct populations of cells in differentiating mTECs (Figure S2C and S2D).

To further characterize *Mycl* expression in intermediate cells, we applied a combination of RNA-FISH and immunofluorescence to mTECs. We examined the expression of *Mycl* relative to FOXJ1 and a marker of secretory cells, *Pigr* (*Polymeric immunoglobulin receptor*, Blackburn et al., 2022) in differentiating mTECs (Figure S2E). Simultaneous visualization revealed that a subset of cells expressed only *Mycl* (11%, Figure S2F and S2G, cyan arrow) and another subset expressed FOXJ1 (20%, Figure S2F and S2G, yellow arrow). Some FOXJ1-expressing cells co-expressed *Mycl* but no *Pigr-*expressing secretory cells expressed *Mycl* (Figure S2F and S2G, magenta arrow). Thus, these data are consistent with the scRNA-seq predictions that *Mycl* expression precedes and overlaps with that of FOXJ1 early in the multiciliated cell lineage.

To assess whether *Mycl* is expressed in intermediate cells *in vivo*, we performed RNA-FISH on embryonic day 17.5 (E17.5) mouse trachea. We multiplexed probes for *Mycl* with those for *Trp63*, *Scgb1a1* and *Foxj1* to label basal stem cells, secretory cells and multiciliated cells, respectively. *Mycl*-expressing cells were abundant in the developing trachea (Figure S2H, cyan arrow) and were mutually exclusive with *Trp63-* or *Scgb1a1*-expressing cells. Some *Mycl*-expressing cells also expressed *Foxj1*, while others did not. Therefore, select cells in the developing mouse trachea express *Mycl*, and *Mycl* expression may precede or be concurrent with markers of multiciliated cells.

### *Mycl* is co-expressed with genes encoding regulators of early multiciliated cell differentiation, *Gmnc* and *Mcidas*

Previous studies, corroborated by our scRNA-seq analysis (Figure 2A), indicated that transcriptional regulators *Gmnc* and *Mcidas* are expressed early in the multiciliated cell lineage (Arbi et al., 2016; Stubbs et al., 2012; Terré et al., 2016). To test whether *Mycl* is also expressed early in the multiciliated lineage, we assessed *Mycl* co-expression with *Gmnc* and *Mcidas*. We performed multiplexed RNA-FISH for *Mycl*, *Gmnc* and *Mcidas* in mTECs cultured for three days at air/liquid interface. At this early stage of differentiation, most *Gmnc*-expressing cells (55%) did not express *Mycl* (Figure 2B, yellow arrow and S3A). However, most *Mycl*-expressing cells (55%) also expressed *Gmnc* (Figure 2B, cyan arrow and S3B), suggesting that *Gmnc* expression is induced in multiciliated cell precursors prior to or independently of *Mycl*. In contrast, most *Mcidas*-expressing cells (77%) also expressed *Gmnc* and *Mycl* or *Mycl* alone (Figure 2B and S3C), further suggesting that *Mcidas* is expressed later than *Gmnc* or *Mycl* in deuterosomal cells of the multiciliated cell lineage.

**Figure 2.**
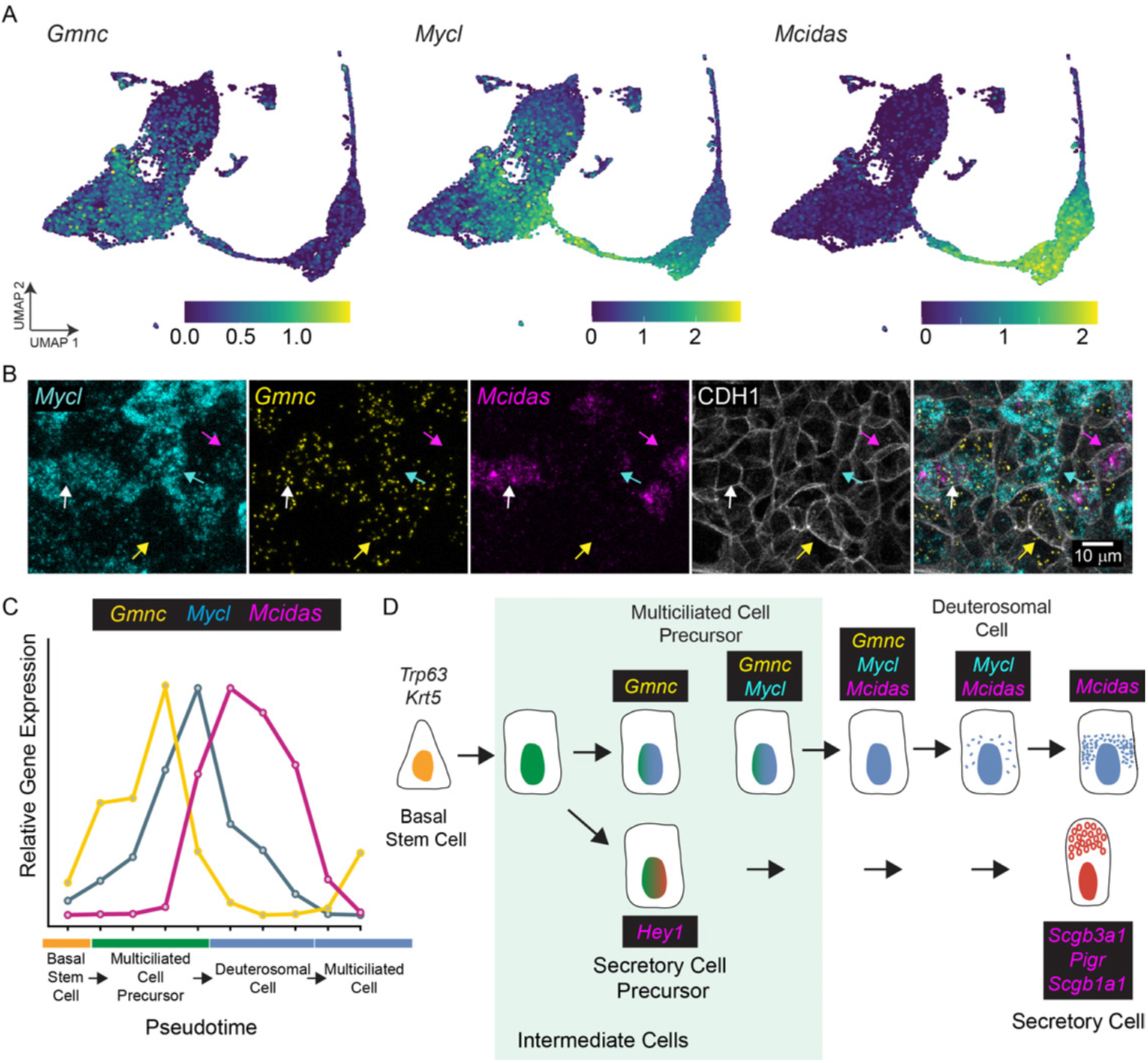
*Mycl* is co-expressed with genes encoding regulators of early multiciliated cell differentiation, *Gmnc* and *Mcidas*. A) Expression of *Gmnc*, *Mycl*, and *Mcidas* superimposed on the UMAP. B) Visualization of *Mycl* transcripts with transcripts of genes expressed by early multiciliated lineage cells (*Gmnc*) and deuterosomal cells (*Mcidas*) in mTECs differentiated for 3 days at air/liquid interface. Cells expressed *Gmnc* only (yellow arrow), both *Mycl* and *Gmnc* (cyan arrow), *Mycl, Gmnc* and *Mcidas* (white arrow), or *Mycl* and *Mcidas* (magenta arrow). C) Relative expression of genes encoding three transcriptional regulators over the pseudotime trajectory from basal stem cells to mature multiciliated cells. *Gmnc* is expressed first, followed by *Mycl* and then *Mcidas*. Expression data is from scRNA-seq of differentiating primary airway epithelial cells (described in Figure 1). D) Schematic of gene expression during basal stem cell differentiation into multiciliated lineage and secretory cells. In the multiciliated cell trajectory, *Gmnc* and then *Mycl* expression initiates in the multiciliated precursors within the intermediate cell population. When producing deuterosomes for basal body synthesis, these cells turn on *Mcidas* and switch off *Gmnc* and *Mycl*. *Hey1* expression initiates in the secretory precursors within the intermediate cell population, then turn on *Scgb3a1*, *Pigr* and *Scgb1a1*.

Together with our scRNA-seq pseudotime analysis, these RNA-FISH analyses support a hierarchy of regulatory expression, with *Gmnc* and *Mycl* expression induced in multiciliated cell precursors, followed by *Mcidas* during early multiciliated cell differentiation (Figure 2C and 2D). These three transcriptional regulators are expressed transiently and in a staggered fashion during differentiation (Figure 2C and 2D).

### GMNC and MCIDAS are required for multiciliated cell differentiation, but not their induction

Based on the sequential expression of *Gmnc* and *Mcidas* in intermediate cells and deuterosomal cells, respectively, we hypothesized that *Gmnc* is required for multiciliated cell fate induction, while *Mcidas* is required for multiciliated cell differentiation. To test these hypotheses, we used CRISPR/Cas9 to disrupt *Gmnc* or *Mcidas* in mTECs. Genomic DNA sequencing revealed that CRISPR/Cas9 efficiently disrupted *Gmnc* and *Mcidas* (referred to as *Gmnc* KO or *Mcidas* KO hereafter) (Figure S4A). Immunofluorescence of *Gmnc* KO or *Mcidas* KO cells differentiated at air/liquid interface for 21 days revealed that, similar to *Gmnc* and *Mcidas* KO phenotypes *in vivo* (Arbi et al., 2016; Lu et al., 2019; Terré et al., 2016), both genes are required for basal body duplication and multiciliogenesis (Figure 3A and S4B). Knockout of either *Gmnc* or *Mcidas* in mTECs decreased expression of MYB and FOXJ1, indicating that *Gmnc* and *Mcidas* are required for early steps of multiciliated cell differentiation (Figure S4C and S4D).

**Figure 3.**
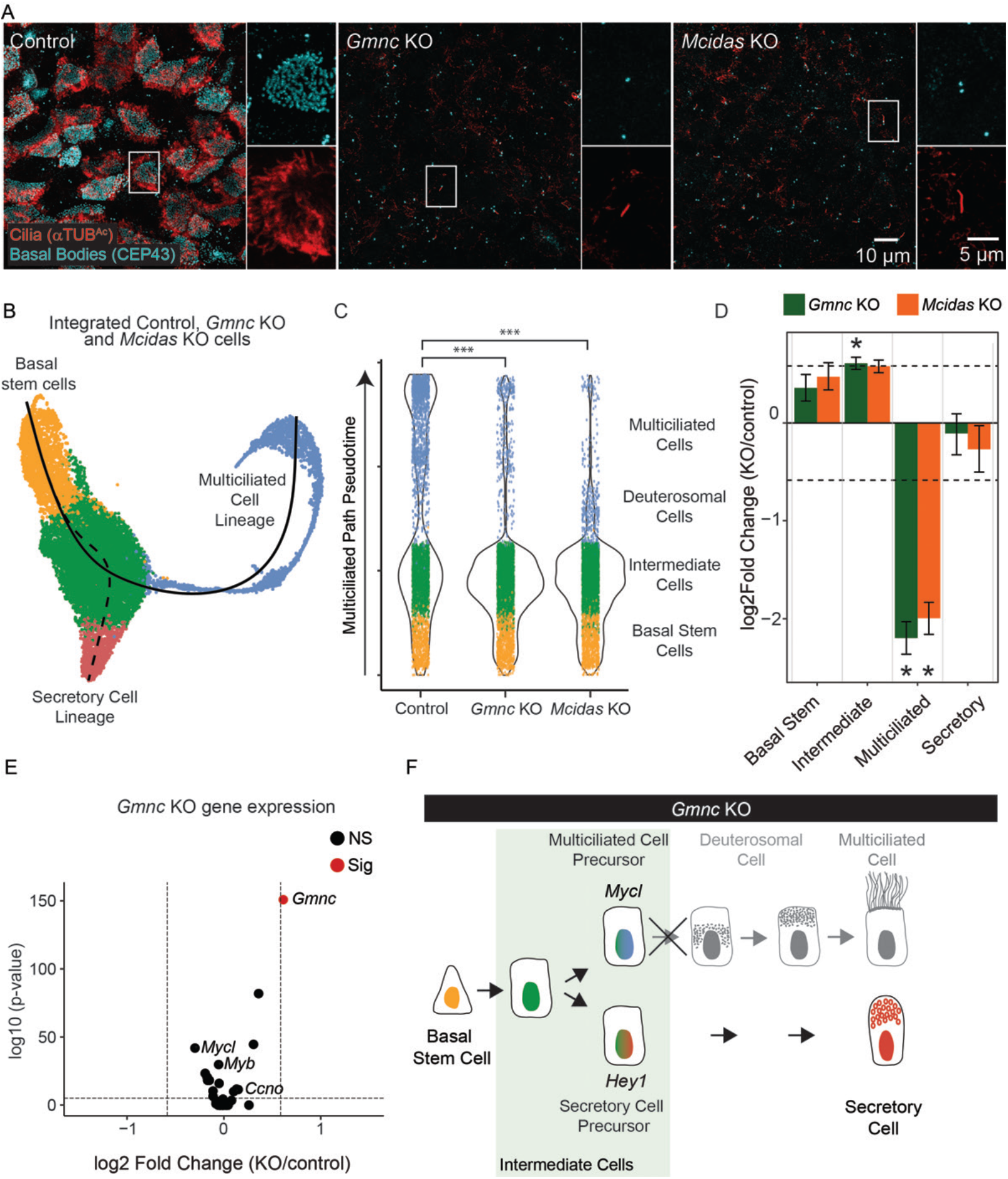
*Gmnc* and *Mcidas* are dispensable for the initiation of the multiciliated lineage. A) Immunofluorescence images of control mTECs (receiving a non-targeting gRNA), *Gmnc* KO mTECs (receiving a gRNA targeting *Gmnc*) and *Mcidas* KO mTECs (receiving a gRNA targeting *Mcidas*). After 21 days at air/liquid interface, cells were immunostained for cilia (acetylated α-tubulin, αTUB^AC^, red) and basal bodies (CEP43, cyan). Insets depict characteristic multiciliated or monociliated cells. B) UMAP, clustering, and Slingshot pseudotime analysis of integrated control, *Gmnc* KO and *Mcidas* KO data. C) Violin plot of pseudotime values along the multiciliated differentiation path for each cell. Cells are colored by clusters as depicted in (B). ***Differential progressionTest (condiments R package) p-value < 0.005. D) Permutation test of cell cluster proportions for *Gmnc* KO and *Mcidas* KO as compared to control. Dotted line demarcates 1.5-fold change. Difference in cluster proportions were considered significant if greater than 1.5-fold change relative to control and FDR < 0.05. Error bars represent 95% confidence interval. E) Volcano plot of *Mycl*-correlated genes in *Gmnc* KO intermediate cells vs. control intermediate cells. Dotted line demarcates 1.5-fold change. Genes with greater than 1.5-fold change in *Gmnc* KO intermediate cells vs. control intermediate cells are highlighted in red. NS=not significant. Sig=significant (Fold-change > 1.5 and p-value < 10e^−6^). F) Model of the effect of loss of *Gmnc* on basal cell differentiation. Without *Gmnc*, intermediate cells accumulate and fail to progress to deuterosomal or mature multiciliated cell stages. *Gmnc* KO intermediate cells maintain expression of *Mycl* and other multiciliated cell precursor genes, suggesting they are specified but unable to differentiate.

To reveal how GMNC and MCIDAS direct multiciliated cell differentiation, we used scRNA-seq to compare the transcriptomes and trajectories of control, *Gmnc* KO and *Mcidas* KO mTECs during early differentiation (*i.e*., after five days of culture at air/liquid interface, Figure S4E and Supplementary Table 4). As in our previous scRNA-seq dataset, pseudotime analysis of integrated control, *Gmnc* KO and *Mcidas* KO scRNA-seq data revealed a differentiation trajectory originating in the basal stem cell cluster, splitting within the intermediate cell cluster and proceeding through either the multiciliated or secretory cell clusters (Figure 3B).

We investigated whether *Gmnc* KO and *Mcidas* KO altered cell states along these trajectories using a differential progression test (Bézieux et al., 2021). Loss of GMNC or MCIDAS significantly altered the distribution of cell states along the multiciliated pseudotime pathway (Figure 3C). To pinpoint where along the multiciliated trajectory this disruption is located, we used permutation testing of cell type proportions (Figure 3D). Neither secretory nor basal stem cell proportions were altered in either knockout (Figure 3D and S4F). Loss of GMNC led to a significant increase in intermediate cells (1.5-fold, FDR = 0.002) and a decrease in multiciliated lineage cells, representing a disruption in the transition from intermediate cell to multiciliated cell differentiation (Figure 3D). Loss of MCIDAS led to an increase in deuterosomal cells of the multiciliated cell lineage and a decrease in mature multiciliated cells, representing a disruption in the progression along the multiciliated cell trajectory (Figure 3C and S4G).

To test whether GMNC is required to initiate multiciliated cell differentiation, we assessed whether GMNC is essential for induction of the multiciliated cell precursor gene signature defined above (Figure 1E). *Gmnc* KO did not alter multiciliated cell precursor gene expression, including *Mycl* expression, within intermediate cells (Figure 3E). Thus, GMNC is dispensable for induction of multiciliated cell precursors and is required instead for their subsequent differentiation into deuterosomal cells (Figure 3F).

### MYCL promotes and HEY1 inhibits multiciliated cell fate

As *Mycl* is expressed by multiciliated precursors within the intermediate cell population, we investigated whether MYCL is sufficient to drive the multiciliated cell fate. We transduced mTECs with no lentivirus, a control lentivirus (expressing a fusion of a nuclear localization signal and GFP, NLS-GFP) or a *Mycl*-expressing lentivirus (MYCL-GFP). To assess whether MYCL affects multiciliated cell differentiation, we analyzed transduced mTECs over the course of differentiation (*i.e*., at 0, 1, 2 and 5 days after transfer to air/liquid interface). Immunofluorescence imaging of cilia and basal bodies revealed that MYCL-GFP expression drove a striking increase in the proportion of multiciliated cells (Figure 4A and 4B), indicating that MYCL expression promotes multiciliated cell differentiation.

**Figure 4.**
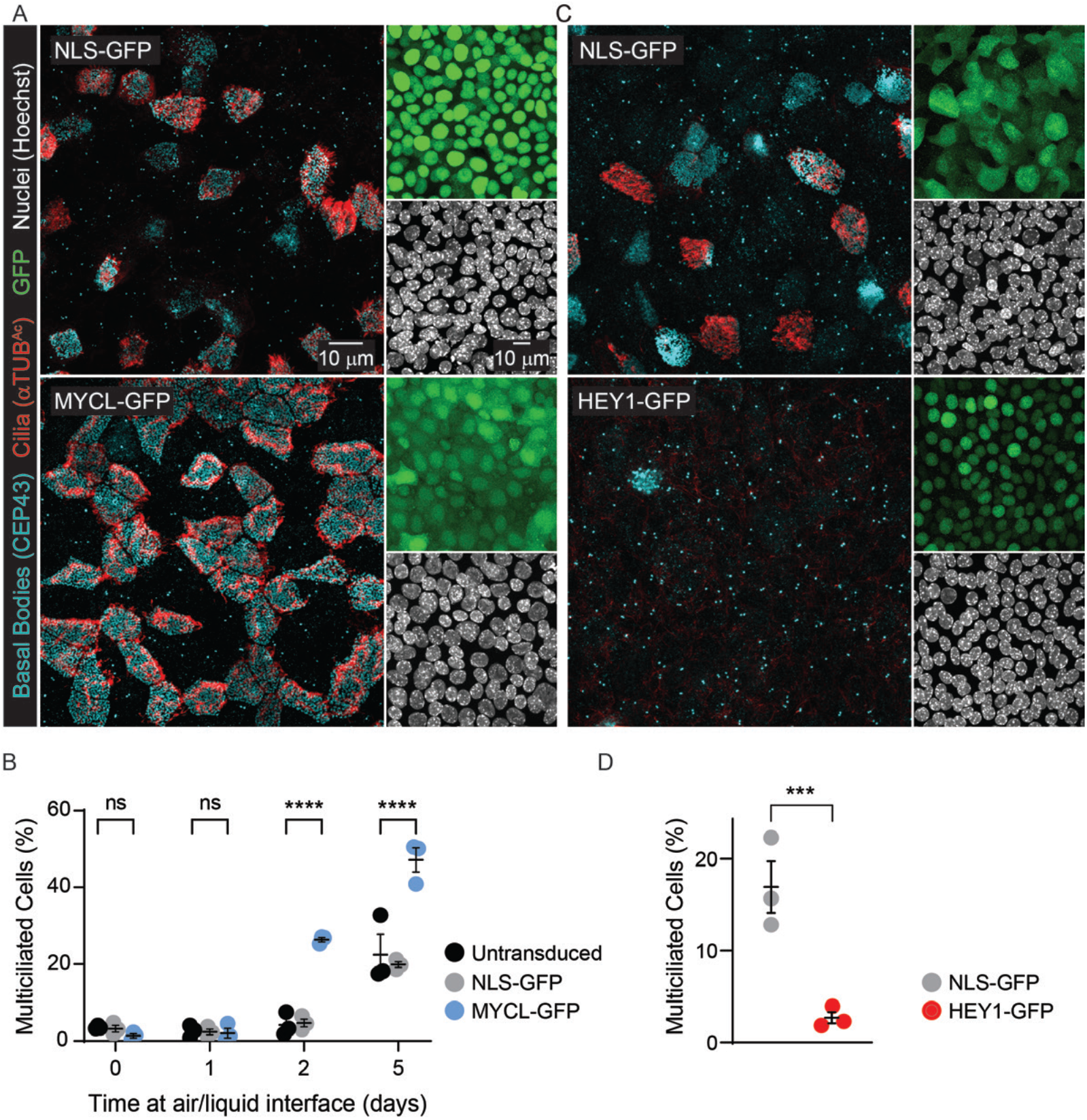
MYCL promotes differentiation of multiciliated cells. A) Immunofluorescence images of mTECs transduced with lentivirus encoding NLS-GFP (control) or MYCL-GFP. Cells were fixed and stained for cilia (αTUB^AC^, red) and basal bodies (CEP43, cyan) five days after differentiation at air/liquid interface. Insets display GFP fluorescence (green) and nuclei (Hoechst, grey). B) Quantification of the percentage of multiciliated cells in untransduced mTEC cultures (black) or mTEC cultures ectopically expressing NLS-GFP (grey) or MYCL-GFP (blue). Multiciliated cell percentage for each condition was counted at several timepoints after initiation of differentiation. Significance was assessed using a two-way ANOVA test with multiple comparison correction, with **** indicating p < 0.0001. Error bars represent SEM for 3 replicates of independently derived and transduced mTECs. C) Immunofluorescence images of mTECs transduced with lentivirus encoding NLS-GFP (control) or HEY1-GFP. Cells were fixed and stained for cilia (αTUB^AC^, red) and basal bodies (CEP43, cyan) five days after differentiation at air/liquid interface. Insets display GFP fluorescence (green) and nuclei (Hoechst, grey). D) Quantification of the percentage of multiciliated cells in mTEC cultures ectopically expressing NLS-GFP (grey) or HEY1-GFP (red). Multiciliated cell percentage for each condition was counted at five days after initiation of air/liquid interface. Significance was assessed using a one-way ANOVA test with multiple comparison correction, with *** indicating p = 0.0003. Error bars represent SEM for 3 replicates of independently derived and transduced mTECs.

As *Hey1* is expressed in secretory cell precursors within the intermediate cell population, and as HEY1 is a transcriptional repressor, we hypothesized that HEY1 could inhibit multiciliated cell fate. To test this hypothesis, we transduced mTECs with a control lentivirus (NLS-GFP) or a *Hey1*-expressing lentivirus (HEY1-GFP). We analyzed transduced mTECs for multiciliated cell fate by immunofluorescence imaging of cilia and basal bodies. We found that HEY1-GFP expression inhibits differentiation into multiciliated cells (Figure 4C and 4D). As *Hey1* is expressed by secretory precursors, we propose that HEY1 helps to inhibit multiciliated cell gene expression in secretory cell precursors within intermediate cells.

### Notch signaling in intermediate cells directs secretory and multiciliated cell fates

In the airway, Notch signaling promotes the secretory cell fate and Notch inactivity promotes the multiciliated cell fate (Rock et al., 2011; Tsao et al., 2009). We hypothesized that Notch signaling acts on intermediate cells to partition them into either *Mycl*-expressing or *Hey1*-expressing lineages.

To test this hypothesis, we activated the Notch signaling pathway in mTECs by expressing GFP-tagged *Notch1* intracellular domain (GFP-N1ICD), a constitutively active Notch effector that inhibits multiciliated cell formation in murine and human airways (Gomi et al., 2015; Guseh et al., 2009). RT-qPCR early after initiation of differentiation revealed that Notch activation inhibited expression of all three early regulators of multiciliated cell fate, *Gmnc, Mycl* and *Mcidas* (Figure 5A).

**Figure 5.**
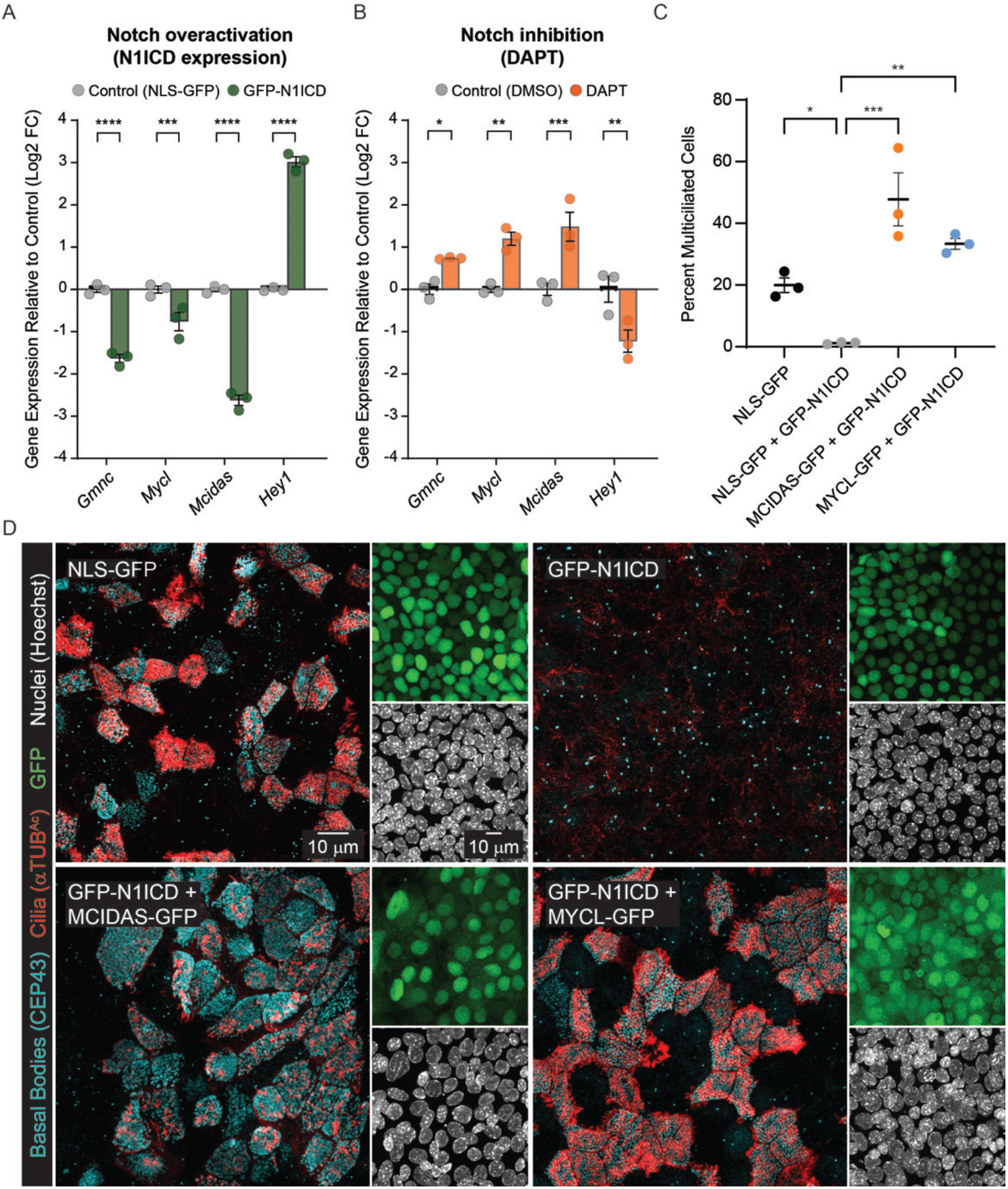
Notch signaling induces *Hey1* and represses *Mycl*. A) qRT-PCR measurement of *Gmnc*, *Mycl*, *Mcidas* and *Hey1* expression in NLS-GFP (control) or GFP-N1ICD transduced mTECs after one day at air/liquid interface. Significance was assessed using a two-way ANOVA test with multiple comparison correction, with **** indicating p < 0.0001 and *** indicating p = 0.0002. Error bars represent SEM for 3 replicates of independently derived and transduced mTECs. B) qRT-PCR measurement of *Gmnc*, *Mycl*, *Mcidas* and *Hey1* expression in mTECs treated with DMSO (control) or a chemical inhibitor of Notch signaling (DAPT) after one day at air/liquid. Significance was assessed using a two-way ANOVA test with multiple comparison correction, with * indicating p = 0.0237, ** indicating p = 0.0021 and *** indicating p = 0.0005. Error bars represent SEM for 3 replicates of independently derived and treated mTECs. C) Quantification of the percentage of multiciliated cells in mTECs ectopically expressing NLS-GFP, GFP-N1ICD, GFP-N1ICD+MCIDAS-GFP or GFP-N1ICD+MYCL-GFP after five days at air/liquid interface. Significance was assessed using a one-way ANOVA test with multiple comparison correction, with * indicating p = 0.0477, *** indicating p = 0.0003 and ** indicating p = 0.0027. Error bars represent SEM for 3 replicates of independently derived and transduced mTECs. D) Immunofluorescence images of mTECs transduced with lentivirus encoding NLS-GFP (control), a Notch-activating protein (GFP-N1ICD) or a Notch-activating protein with either MCIDAS (GFP-N1ICD+MCIDAS-GFP) or MYCL (GFP-N1ICD+MYCL-GFP). After five days at air/liquid interface, cells were stained for cilia (aTUB^AC^, red), basal bodies (CEP43, cyan) and nuclei (Hoechst, gray). GFP fluorescence (green) demonstrates transgene expression.

Conversely, we inactivated Notch signaling using the g-secretase inhibitor, DAPT, which strongly induces multiciliated cell formation (Gerovac et al., 2014; Gomi et al., 2015; Vladar et al., 2016). Notch inhibition induced *Gmnc, Mycl* and *Mcidas* expression (Figure 5B). Together, these experiments demonstrate that *Mycl*, like *Gmnc* and *Mcidas* (Kyrousi et al., 2015; Stubbs et al., 2012; Zhou et al., 2015), is repressed by Notch signaling.

Notch inactivation could increase *Mycl* expression by increasing the proportion of *Mycl*-expressing intermediate cells or the *Mycl*-expressing early multiciliated cells. As intermediate cells do not express FOXJ1 but early multiciliated cells do, we combined multiplexed RNA-FISH and immunofluorescence to simultaneously visualize expression of *Mycl* and FOXJ1 in GFP-N1ICD-transduced cells. Notch activation reduced the proportion of both *Mycl*-expressing intermediate cells (*i.e*., cells expressing *Mycl* but not FOXJ1) and early multiciliated cells (*i.e*., cells expressing *Mycl* and FOXJ1, Figure S5A and S5C). Conversely, Notch pathway inhibition with DAPT increased the proportion of *Mycl*-expressing intermediate cells early in differentiation (one day after transfer to air/liquid interface) (Figure S5A and S5C) leading to a dramatic increase in the number of multiciliated cells two days later (Figure S5B and S5D). Thus, Notch signaling acts upstream of *Mycl* within the intermediate population to direct cells to become either multiciliated or secretory cell precursors.

We reasoned that if MYCL acts downstream of Notch signaling, MYCL could drive multiciliated cell fate even in the presence of active Notch signaling. To test this prediction, we simultaneously expressed GFP-N1ICD together with NLS-GFP (control), MCIDAS-GFP or MYCL-GFP (Figure 5C). As expected, GFP-N1ICD suppressed formation of multiciliated cells. MCIDAS-GFP restored multiciliated cell formation to mTECs with activated Notch (Figure 5C and 5D), consistent with epistasis data from the frog epidermis (Stubbs et al., 2012). MYCL-GFP also restored multiciliated cell formation in the presence of activated Notch (Figure 5C and 5D). Thus, MYCL is epistatic to Notch, and Notch signaling represses multiciliated cell fate, at least in part, by repressing *Mycl* expression in intermediate cells.

### MYCL and HEY1 bind the same regulatory elements near multiciliated genes to direct basal stem cell differentiation

To investigate how MYCL and HEY1 promote precursor cell identity, we identified where these transcription factors bind the genome in airway epithelial cells using CUT&RUN (Cleavage Under Targets and Release Using Nuclease, Skene and Henikoff, 2017). Comparison of CUT&RUN data from mTECs expressing NLS-GFP (control), MYCL-GFP or HEY1-GFP identified 13,398 MYCL-GFP peaks and 13,758 HEY1-GFP peaks. Over half of both MYCL and HEY1 peaks were located within 3 kb of gene promoters (Figure 6A). Surprisingly, most (60%) MYCL and HEY1 peaks overlapped, suggesting they may antagonize each other via opposing effects at shared regulatory regions (Figure 6B).

**Figure 6.**
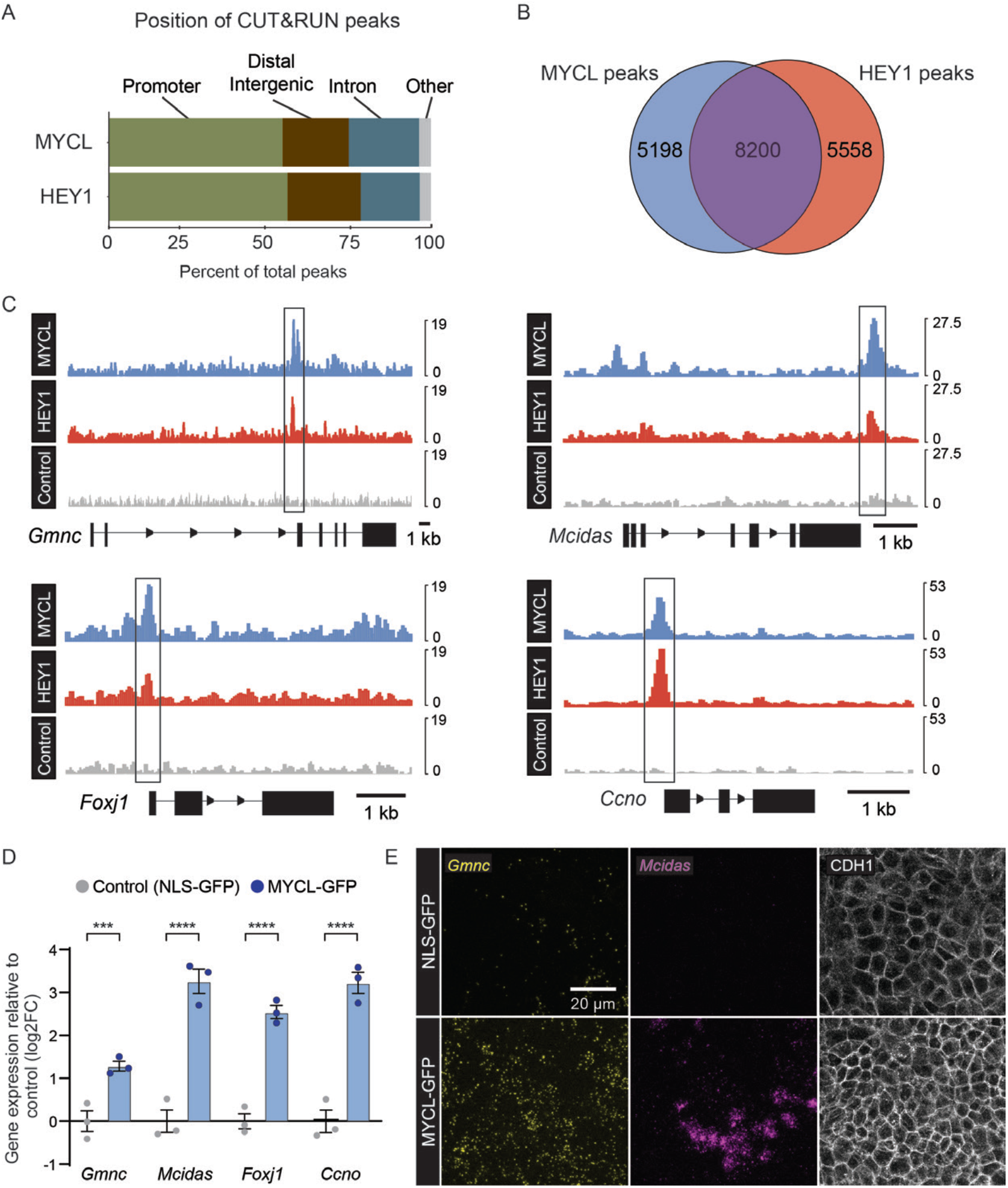
MYCL and HEY1 directly regulate early multiciliated lineage genes. A) mTECs transduced with MYCL-GFP, HEY1-GFP or a control (NLS-GFP) lentivirus were analyzed using CUT&RUN after one day at air/liquid interface. Distribution of MYCL and HEY1 peaks relative to gene positions. B) Venn diagram of the overlap of MYCL and HEY1 binding sites in mTECs. C) MYCL, HEY1 and control CUT&RUN signals near the *Gmnc*, *Mcidas, Foxj1*, and *Ccno* loci. Gene models are depicted 5’ to 3’. Regions with identified peaks are outlined in black. Scale bars depict 1 kilobase pairs of DNA. D) qRT-PCR measurement of *Gmnc*, *Mcidas, Foxj1* and *Ccno* expression in control (NLS-GFP) MYCL-GFP transduced mTECs cultured for one day at air/liquid interface. Significance was assessed using a two-way ANOVA test with multiple comparison correction, with *** indicating p = 0.0009 and **** indicating p < 0.0001. Error bars represent SEM for 3 replicates of independently derived and transduced mTECs. E) Multiplexed RNA-FISH and immunofluorescence of control (NLS-GFP) or MYCL-GFP transduced mTECs cultured for one day at air/liquid. *Gmnc* and *Mcidas* staining was multiplexed with immunostaining for CDH1 to mark cell boundaries.

Identifying the gene nearest to peaks revealed that multiple shared MYCL and HEY1 peaks occur near genes encoding early, critical regulators of the multiciliated cell fate (*i.e*., *Gmnc*, *Mcidas*, *Foxj1 and Ccno*) (Figure 6C and Supplementary Table 5). To test whether MYCL can induce these early ciliogenic genes, we quantitated the effects of MYCL-GFP on mTEC gene expression using RT-qPCR (Figure 6D). MYCL-GFP robustly induced expression of *Gmnc*, *Mcidas, Foxj1* and *Ccno*, indicating that MYCL may directly activate regulators of multiciliated cell differentiation.

To further test whether MYCL activates the early multiciliated cell differentiation program, we performed RNA-FISH to assess *Gmnc* and *Mcidas* expression in mTECs expressing NLS-GFP or MYCL-GFP. Early in differentiation, NLS-GFP-expressing control mTECs included few cells expressing either *Gmnc* or *Mcidas*. In contrast, at the same early point in differentiation, MYCL-GFP-expressing mTECs exhibited dramatically increased numbers of cells expressing both *Gmnc* and *Mcidas* (Figure 6E). Thus, MYCL can accelerate induction of the early multiciliated lineage transcriptional program.

## Discussion

Using single cell gene expression analysis of primary airway epithelial cells, we identified a differentiation trajectory from basal stem cells to multiciliated and secretory cells. This trajectory bifurcates between basal stem, multiciliated and secretory cells in an intermediate population of cells. Notch decides between the multiciliated and secretory cell fates in this intermediate cell state: active Notch induces the transcriptional repressor *Hey1* and drives secretory cell fate while inactive Notch leads to the expression of the transcriptional activators *Gmnc* and *Mycl* and drives multiciliated cell fate. MYCL binds regulatory elements near multiciliated cell genes and induces their expression, thereby promoting multiciliated cell differentiation. HEY1 binds many of the same regulatory elements and represses expression of the multiciliated cell program in secretory cell precursors. Thus, this work identifies two antagonistic effectors downstream of Notch signaling that funnel differentiating basal stem cells into two alternative fates (Figure 7).

**Figure 7.**
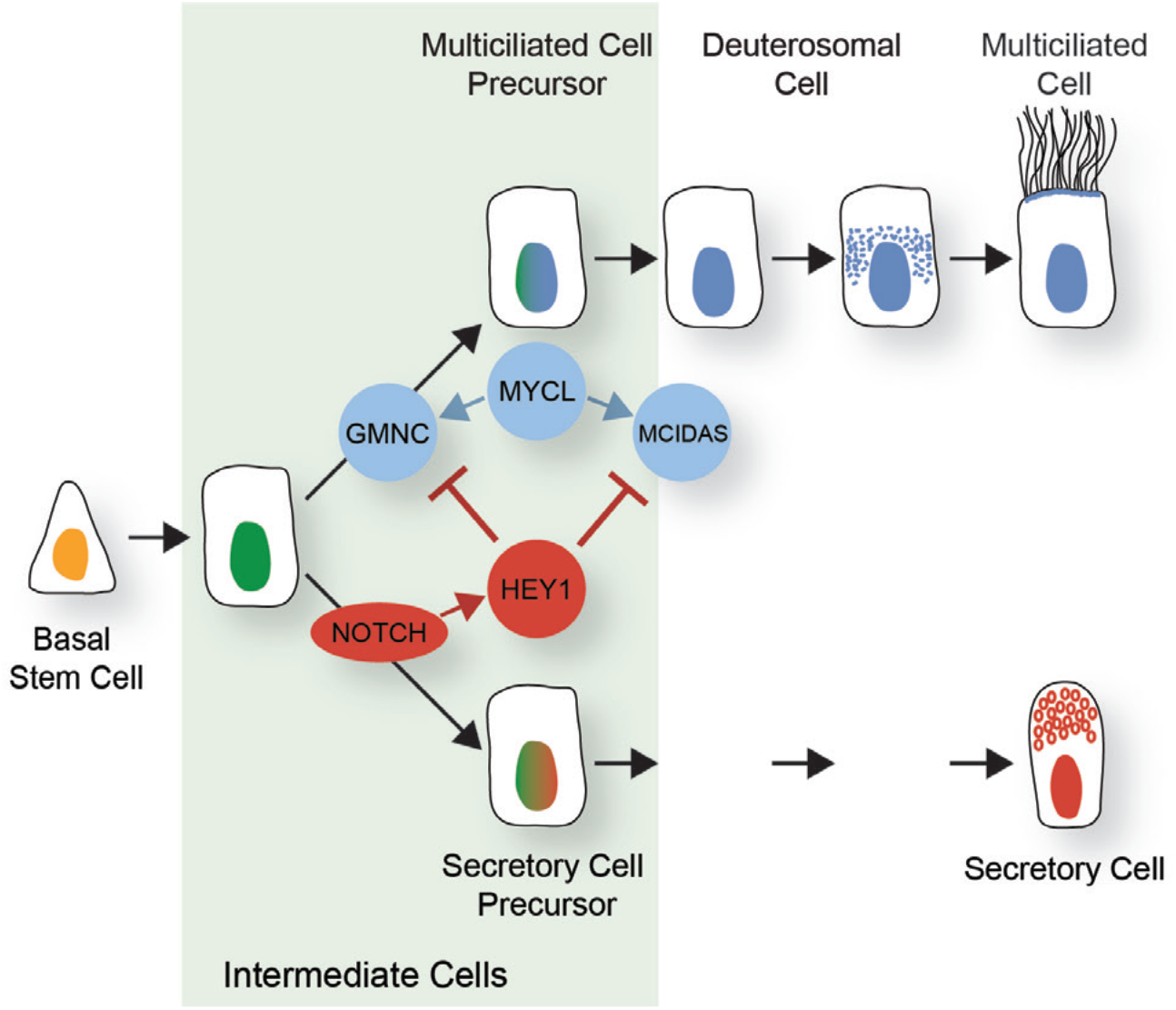
A model for the differentiation of airway stem cells. Schematic model of the Notch-dependent cell fate decision that takes place in intermediate cells during basal stem cell differentiation. Intermediate cells that do not see active Notch signaling turn on a network of GMNC, MYCL and MCIDAS to become multiciliated cell precursors. Intermediate cells that activate the Notch pathway switch on HEY1 to repress the multiciliated cell gene expression program and acquire a secretory cell precursor identity.

A population of airway epithelial cells which, akin to intermediate cells, lack markers of basal stem cells and differentiated cells has been previously described and, reflecting their location in the epithelium, were termed suprabasal or parabasal cells (Donnelly et al., 1982; Mercer et al., 1994). It was proposed that these suprabasal cells derive from basal stem cells and give rise to either secretory or multiciliated cells (Mori et al., 2015; Rock et al., 2011; Watson et al., 2015). *Notch3* and *Krt8* expression has been suggested to mark these suprabasal cells (Mori et al., 2015; Rock et al., 2011; Watson et al., 2015). We find that the intermediate cells described in this work express both *Notch3* and *Krt8* at levels higher than basal stem cells (Figure S6A), supporting the idea that airway suprabasal cells may be in the intermediate cell state.

Lineage tracing of *Scgb1a1*-expressing cells previously suggested that secretory cells can differentiate into mature multiciliated cells (Rawlins et al., 2009). We observed low levels of *Scgb1a1* expression in intermediate cells (Figure S6B), which could be sufficient for Cre-dependent activation. Our data are consistent with the possibility that intermediate cells and previously described club cells are the same population, with cells in both categories able to give rise to mature secretory and mature multiciliated cells. The *Scgb1a1* expression in intermediate cells could also explain the differences in cell type labeling observed in *Scgb1a1-Cre* transgenic mice (Kathiriya et al., 2020; Rawlins et al., 2009).

After injury, Notch signaling acts on basal stem cells to direct them to directly become multiciliated lineage cells or secretory lineage cells (Pardo-Saganta et al., 2015). As we did not identify distinct populations of basal stem cells among mTECs, it is likely that Notch acts differently during injury repair and in primary cultured tracheal cell differentiation, the former using direct differentiation and the latter a Notch-mediated decision between two fates during the intermediate cell state.

This intermediate cell state is defined by a composite expression profile, rather than any individual marker. Intermediate cells exist at the intersection of transcriptional gradients defining basal stem cells and differentiating cells. While unique markers of the intermediate cell state may exist and simply be undetectable by scRNA-seq, the absence of unique markers is consistent with the possibility that the intermediate cell state is transitory, present after the downregulation of the basal stem cell program and before the upregulation of either the secretory or multiciliated cell programs. Whether other types of differentiating stem cells also pass through a similar intermediate state defined by a nadir of state-specific transcription will be interesting to assess. One possibility is that erasing the stem cell-specific transcriptional program generates a relatively blank slate on which new directions for lineage commitment can be written.

While later stages of multiciliated cell differentiation have been described by scRNA-seq (Deprez et al., 2020; García et al., 2019; Montoro et al., 2018; Plasschaert et al., 2018), here we identified the earliest stage of the multiciliated cell lineage. We found that multiciliated cell precursors in the intermediate state express a transient pulse of *Gmnc* and *Mycl* to promote adoption of the multiciliated cell lineage. The finding that *Mycl* expression in these precursors is independent of *Gmnc* suggests that, surprisingly, *Gmnc* is not required for the induction of multiciliated precursors but rather for their differentiation. The pulse of GMNC and MYCL then induces a pulse of *Mcidas* to propel the precursors to become deuterosomal cells. The ability of MYCL to directly induce the expression of *Gmnc* and *Mcidas* and drive multiciliated cell differentiation (even in the presence of active Notch signaling) suggests that GMNC and MYCL cooperatively direct intermediate cells to adopt the multiciliated lineage.

We also found that the transcriptional repressor HEY1 is induced among intermediate cells to oppose MYCL and promote adoption of the secretory lineage. HEY1 is induced by Notch signaling and binds to regulatory regions near genes involved in early multiciliated cell differentiation. We propose that HEY1 represses the early multiciliated cell transcriptional program in secretory cell precursors, thereby reinforcing the secretory cell fate decision downstream of Notch activation. Consistent with this model, HEY1 misexpression reduced multiciliated cell differentiation. Discovering whether Notch signaling has additional roles at later points of cell differentiation awaits further investigation. However, the ability of Notch inhibition to change the balance of multiciliated and secretory cells in adult lung suggests that Notch signaling may also be required to maintain secretory cell differentiation (Lafkas et al., 2015).

The unexpected finding that MYCL and HEY1 bind many of the same regulatory regions suggests that these factors antagonistically regulate the multiciliated cell precursor expression program in intermediate cells. Both MYCL and HEY1 have previously been shown to interact with canonical and variant E-boxes (Amati et al., 1992; Blackwood and Eisenman, 1991; Fischer et al., 2002; Heisig et al., 2012; Mukherjee et al., 1992). Depending on Notch activity in each intermediate cell, only one of these two bHLH transcriptional regulators may bind regulatory elements near multiciliogenic genes, with MYCL promoting and HEY1 inhibiting the adoption of multiciliated cell fate.

Altered proportions of cell types is a hallmark of diverse diseases. For example, blood dyscrasias are caused by an overabundance (*e.g*., erythrocytosis) or lack (*e.g*., anemia) of constituent cell types. Airway diseases are similarly associated with altered cell type proportions: hyperplasia of secretory cells at the expense of multiciliated cells is a hallmark of chronic asthma and COPD. One mechanism by which cell type proportions can be disrupted is via altered Notch signaling (Hartenstein and Posakony, 1990; Heitzler and Simpson, 1991). Indeed, inhibiting Notch signaling in the airway can restore homeostatic proportions of cell types (Lafkas et al., 2015), underscoring that understanding the molecular mechanisms by which Notch signaling directs these two fates is important for understanding airway development, the etiology of lung pathology, and the effectors of potential therapeutic approaches. Here, we have identified how Notch acts through two opposing transcriptional effectors, MYCL and HEY1, to balance the formation of two alternative fates. We propose that dysregulation of these factors may contribute to the imbalance of secretory and multiciliated cells in disease, and approaches leading to activation of MYCL or inhibition of HEY1 may allow for restoration of multiciliated cells and clearance of overabundant mucus in airway disease.

### Limitations of the study

We have shown that overexpression of two novel transcriptional regulators of airway epithelial fate, MYCL and HEY1, can drive multiciliated cell fate or block multiciliated cell fate, respectively. Our pseudotime analysis predicts that intermediate cells expressing *Mycl* or *Gmnc* become multiciliated cells, and that intermediate cells expressing *Hey1* become secretory cells. Validation of these computational predictions will require the development of tools to lineage trace these subpopulations of intermediate cells. Complementary approaches, such as bulk RNA-sequencing or mass spectrometry of isolated intermediate cells, may help to identify additional novel markers of these intermediate cells, allowing us to develop molecular reagents to further define their differentiation potential and lineage commitment. Testing the hypothesis emerging from this work that MYCL and HEY1 compete for binding at shared regulatory elements will require quantitative assessment of regulatory element occupancy in cells co-expressing various levels of both transcription factors.

## Materials and Methods

### Mouse husbandry

Wild-type adult mice of the strain *C57BL/6J* were obtained from Jackson Labs for these studies (JAX Stock #000664). *Foxj-CreERT* (*Foxj1*^tm1.1(cre/ERT2/GFP)Htg^) mice were obtained from the Jackson Laboratory (Stock #027012) and have been described previously (Muthusamy et al., 2014). *Rosa-tdTomato* (*Gt(ROSA)*^26Sortm14(CAG-^ tdTomato)Hze) mice were obtained from the Jackson Laboratory (JAX Stock #007914) and have been described previously (Madisen et al., 2010). Mice were maintained under standard pathogen-free conditions at the UCSF CVRI animal care facility. All animal protocols and procedures were approved by the Institutional Animal Care and Use Committee (IACUC) of the University of California San Francisco.

### Isolation and culturing of mouse tracheal epithelial cells

Mouse tracheal epithelial cells (mTECs) were purified and cultured as previously described (You and Brody, 2013). Briefly, adult (2-4 months old) mice were anesthetized with isoflurane. Trachea were isolated and placed in 3 mg/ml Pronase (Sigma-Aldrich, Cat #10165921001) in Ham’s F-12 medium supplemented with 100 U/mL penicillin and 100 mg/mL streptomycin. After overnight incubation at 4°C, trachea were mechanically agitated, cells were collected and plated on Primaria plates (Corning, #087724A) in mTEC Basic medium (DMEM/F-12 with 15 mM HEPES, 4 mM L-glutamine, 3.6 mM NaHCO3, 100 U/mL penicillin and 100 mg/mL streptomycin) supplemented with 10% FBS for 4 hours at 37°C to remove contaminating fibroblasts. Epithelial cells in suspension were collected in mTEC Plus medium [mTEC Basic supplemented with 10 μg/ml insulin (Sigma-Aldrich, Cat #I1882), 5 μg/ml transferrin (Sigma-Aldrich, Cat #T1147), 0.1 μg/ml cholera toxin (Sigma-Aldrich, Cat #C8052), 25 ng/ml epidermal growth factor (Corning, Cat #CB40001), 30 μg/ml bovine pituitary extract (Sigma-Aldrich, Cat # P1476), 5% FBS and 50 nM Retinoic Acid (Sigma-Aldrich, Cat # R2625) and 10 μM Rock inhibitor (Y-27632, Sigma-Aldrich, Cat #Y0503)].

mTECs were seeded on 6.5 mm transwells (Corning, Cat #3470) coated with 50 μg/ml rat tail collagen (Type I, Corning, Cat #CB354249) at a density of 33,000 cells per well. mTECs were allowed to proliferate for five days before initiating differentiation by removing media from the apical chamber and changing media in the basal chamber to mTEC Basic with 2% Nu-Serum (Corning, Cat #355100) and 50 nM retinoic acid. Placement at air/liquid interface was considered day zero of differentiation. Subsequent to placement at air/liquid interface, basal media was changed and the apical surface was rinsed twice with PBS every other day.

### CRISPR/Cas9-mediated gene knockout in mTECs

mTECs were plated on collagen-coated 24 mm transwells (Corning, Cat #3450) at a density of 175,000 cells per well in mTEC Plus Expansion medium [mTEC Plus supplemented with 50 nM retinoic acid, 10 μM ROCK inhibitor, 1 μM A83-01 (Tocris, Cat #2939), 200 nM DMH-1 (Tocris Cat #4126) and 500 nM CHIR99021 (Tocris, Cat #4423)]. mTEC Plus Expansion medium formulation was modified from previously described medium (Mou et al., Cell Stem Cell, 2016). Cells were expanded with daily medium changes for 4 days and then removed from the transwell using 0.05% trypsin diluted in Cell Dissociation Buffer (Gibco, Cat #13151014). Cells were collected, counted, washed in PBS, then resuspended in P3 electroporation buffer (Lonza, Cat # V4SP-3096) at a concentration of 20,000 cells per 1 μl.

Guide RNAs targeting *Gmnc*, *Mcidas* or a control nontargeting guide were designed and synthesized (Synthego) (see Resource Table). RNPs were assembled by adding 90 pmol TrueCut Cas9 v2 protein (Thermo Fisher, Cat #A36499) to 180 pmol of sgRNA and incubating at room temperature for 15 min. 400,000 cells in 20 μl of P3 buffer were added to each RNP and electroporated using program 96-EA-104 on an Amaxa 4D-Nucleofector with a 96-well shuttle (Lonza # AAF-1003B/S). Immediately after electroporation, cells from each electroporation were collected in mTEC Plus Expansion medium and seeded onto five 6.5 mm collagen-coated transwells. Cells were expanded with daily media changes until they reached confluence (3-4 days post electroporation) at which point differentiation was initiated as described above.

To assess knockout efficiency, genomic DNA was extracted from a single 6.5 mm transwell using the AllPrep DNA/RNA Mini Kit (Qiagen, Cat #80204). Genomic DNA was PCR amplified using oligonucleotides surrounding the sgRNA binding site (see Resource Table). PCR amplicons were purified, and Sanger sequencing was performed using the corresponding forward primer. Sequence chromatograms were analyzed using Inference of CRISPR Edits (ICE) to evaluate gene knockout efficiency (https://ice.synthego.com/). A minimum of three independently derived and electroporated mTEC populationss per guide RNA were analyzed to calculate knockout efficiency. Significance testing was performed in Prism 9 (GraphPad Software), using an ordinary one-way ANOVA test with Dunnett’s multiple hypothesis correction.

### scRNA-seq

To generate single-cell suspensions for scRNA-seq from mTEC cultures, mTECs derived from wild-type mice and *Foxj1-CreERT Rosa-tdTomato* mice treated with 4-OHT on day two at air/liquid interface were cultured and differentiated. On day three at air/liquid interface, mTECs were washed with PBS three times both apically and basally. Trypsin-EDTA at a concentration of 0.05% in Cell Dissociation Buffer (Gibco, Cat #13151014) was added both apically and basally and cells were incubated at 37°C for 10 min. Cells were lifted from the membrane with gentle pipetting and trypsin deactivated in mTEC Basic with 2% Nu-Serum (Corning, Cat #355100). Fresh 0.05% Trypsin-EDTA in Cell Dissociation Buffer was added to the apical surface and incubated at 37°C for 5 min. The pipetting, inactivation, and 5 min incubation steps were repeated twice. Cells were washed in FACS buffer (2% FBS, 2% BSA, 5 mM EDTA, 25 mM HEPES in PBS) filtered through a 100 μM cell strainer and resuspended in FACS buffer with Fixable Viability dye (Invitrogen, Cat #65-0863-14) at a dilution of 1:1600. After a 15 min incubation on ice, cells were washed with FACS buffer and sorted on a fluorescence activated cell sorter (Sony SH800). tdTomato-expressing *Foxj1-CreERT Rosa-tdTomato* cells, representing cells within the multiciliated cell lineage, were sorted into mTEC Basic with 2% Nu-Serum. Wild-type cells and a fraction of *Foxj1-CreERT Rosa-tdTomato* cells were not sorted. Both tdTomato-expressing and unsorted cells were resuspended in 0.04% BSA in PBS at 1000 cells per μL to prepare for scRNA-seq.

10,000 cells from the sorted tdTomato-expressing and unsorted fractions of *Foxj1-CreERT Rosa-tdTomato* mTECs were loaded onto two separate wells of the 10X Chromium Controller using the Chromium Single Cell 3’ Reagent kit (10X Genomics, version 3, Cat #PN-1000075). In addition, 25,000 cells from two independent wild type mTEC populations were loaded on separate days on the 10X Chromium Controller using the Chromium Single Cell 3’ Reagent kit. Also, 16,000 cells of *Gmnc* KO, *Mcidas* KO, or control mTECs at air/liquid interface for five days were loaded on three separate wells of the 10X Chromium Controller using the Chromium Next GEM Single Cell 3’ Reagents kit. Manufacturer’s instructions were followed for Gel Beads-in-emulsion (GEM) generation, cDNA production and library construction. Libraries were sequenced on the Illumina NovaSeq. CellRanger 3.0 was used with default settings to de-multiplex, align reads to the mouse genome (10X Genomics pre-built mm10 reference genome) and count unique molecular identifiers (UMIs). SoupX (Young and Behjati, 2020) was used to remove ambient background RNA from droplets for all individual datasets. Doublets were removed from individual datasets with Scrublet (Wolock et al., 2019), with an assumed doublet rate based on loading concentrations and the 10X Genomics estimates. Seurat versions 3 and (Stuart et al., 2019) were used for downstream analysis of individual datasets, including filtering, dimension reduction, clustering, UMAP, and differential gene expression analysis.

The four individual scRNA-seq datasets (tdTomato-expressing, unsorted, wild type replicate 1, and wild type replicate 2) were merged in Monocle3 (Cao et al., 2019) using Batchelor (Haghverdi et al., 2018) to correct for batch effects between datasets. Dimension reduction, clustering with the Leiden algorithm, and trajectory inference were performed in Monocle3. Monocle3 UMAP dimensions, cluster IDs, and pseudotime values were imported into a Seurat object to analyze differential gene expression and generate gene expression visualizations. Marker genes were identified using FindAllMarkers. Data was compared to previously generated scRNA-seq datas using Seurat’s label transfer tutorial.

*Gmnc* KO, *Mcidas* KO, and control scRNA-seq datasets were integrated following Seurat’s integration tutorial. Scaling, dimension reduction, and clustering were performed in Seurat to integrate data for visualization. Differential gene expression analysis and gene expression visualization were performed in Seurat on the original, non-integrated values, as recommended by the authors of Seurat. Slingshot (Street et al., 2018) was used to infer trajectories within the integrated UMAP. The Condiments package was used to test differential progression along the trajectories (Bézieux et al., 2021). Cluster proportion testing was performed with available scripts (https://github.com/rpolicastro/scProportionTest).

### Lentiviral transduction

Coding sequences of mouse cDNAs were PCR amplified (*Mycl* [ENSMUST00000030407.8], *Hey1* [ENSMUST00000042412.5] and *Mcidas* [ENSMUST00000092089.6]) or synthesized (NLS and N1ICD the fragment of [ENSMUST00000028288.5] encoding the C-terminal 789 AA of the *Notch1* protein) and cloned with sequence encoding an N- or C-terminal eGFP into a modified version of pLKO.3G (Addgene, Cat #14748, with the sgRNA expression cassette removed). Plasmids generated were NLS-GFP-Lenti, MYCL-GFP-Lenti, MCIDAS-Lenti, HEY1-GFP-Lenti and GFP-N1ICD-Lenti. Lentivirus was made by transfecting 7.5 μg of each plasmid along with 1.5 μg of pCMV-VSV-G (Addgene, Cat #8454) and 6 μg of psPAX2 (Addgene, Cat #12260) into a 10 cm plate of HEK293T cells at 60-70% using Fugene 6 transfection reagent (Promega, Cat #E2691). Media containing lentiviral particles were collected at 24, 48 and 72 hours after transfection and concentrated by centrifugation at 25,000 RPM for 1.5 hours at 4°C and the pellet resuspended in 300 μl PBS. Viruses were frozen and stored at −80°C, and virus from a single preparation was used at the same dosage for the remainder of the experiments.

For lentiviral transduction, mTECs were plated in 6.5 mm transwells and transduced with 1-15 μl of lentivirus in mTEC Plus medium supplemented with 50 nM retinoic acid and 10 μM ROCK inhibitor, as modified from a previously transduction protocol (Horani et al., 2013). Basal media was changed 24 hours after transduction and apical media was changed 48 hours after transduction. Media in both chambers was changed at 72 hours post transduction and mTECs were differentiated, fixed and processed for immunofluorescence imaging.

### Notch signaling modulation

Notch signaling was inhibited by adding 1 μM γ-Secretase inhibitor (DAPT, Sigma, #565784) to mTECs at the initiation of differentiation (air/liquid interface day 0). Control cells were treated with vehicle (DMSO). Notch signaling was activated by transducing mTECs with GFP-N1ICD lentivirus or control lentivirus (NLS-GFP) at the time of seeding. RNA was extracted from cells after 1 day at air/liquid interface for gene expression analysis. Each experiment was performed on three independently derived and treated mTEC cultures. For N1ICD-GFP co-expression with NLS-GFP, MYCL-GFP or MCIDAS-GFP, cells were simultaneously transduced with both lentiviruses.

### Immunofluorescence imaging of mouse tracheal epithelial cells

A minimum of three regions of the membrane were imaged per biological replicate for each condition. mTECs on transwells were washed three times in PBS, and then fixed using in 4% PFA in PBS for 15 min at room temperature. Fixative was removed and cells were rinsed three times with PBS. Transwell membranes were removed from the support using a scalpel and then cut into smaller pieces. An individual piece of membrane was placed on a slide inside a hydrophobic boundary for staining. The remaining steps were carried out in a humidity chamber. Cells were blocked for 1 hour at room temperature in PBT (1% BSA, 0.5% TritonX-100 and 0.02% sodium azide in PBS) supplemented with 10% normal donkey serum, then incubated in primary antibody diluted in PBT for 1 hour at room temperature. After five PBS washes of five min each, secondary antibodies, diluted in PBT, were added along with Hoechst to stain nuclei. Cells were incubated in secondary antibody for 1 hour at room temperature, followed by five PBS washes of five min each. Membranes were mounted in Prolong Diamond Antifade (Molecular Probes, Cat #P36970) mountant. Specifics of primary and secondary antibodies are provided in the Resource Table.

Immunofluorescence images were acquired on a Zeiss 800 laser scanning confocal microscope with a 63x oil objective. A minimum of three regions of the membrane were imaged per biological replicate for each condition. Imaging positions were selected randomly, using the nuclear stain to ensure healthy cells were present. For GFP-N1ICD-expressing cell imaging, regions of the membrane with GFP fluorescence reflecting transgene expression were selected. For HEY1-GFP-expressing cell imaging, only regions of the membrane with nuclear GFP fluorescence were selected. At each position, cells were imaged with a 63x/1.4 objective, collecting between 10-20 images per region, spaced 0.5 μm apart in the z-axis. While collecting images, the gain and laser power were held constant for each antibody combination per experiment. Images were processed identically using ImageJ software to generate maximum projections. Counting was performed manually on maximum projection images, with multiciliated cells defined as cells with multiple cilia (>5 αTUB^AC^-positive objects) or multiple basal bodies (>5 CEP43-positive objects) per cell. Cell number was determined by counting nuclei. All images were processed using ImageJ software. Significance testing was performed in Prism 9, using an ordinary one-way ANOVA test with Dunnett’s multiple hypothesis correction. Figures were assembled in Illustrator (Adobe).

### Quantitative reverse transcription and PCR

mTECs on 6.5 mm transwells were washed three times in PBS. Total RNA was extracted using the RNeasy Plus Mini Kit (Qiagen, Cat #74134) or both gDNA and total RNA were extracted using the AllPrep DNA/RNA Mini Kit (Qiagen, Cat #80204). Reverse transcription was carried out for 1 hour at 42°C using the iSCRIPT cDNA synthesis kit (Bio-Rad, Cat #1708841BUN) and 300 ng - 1 μg of RNA per reaction. Quantitative PCR was performed on a QuantStudio 5 real-time PCR machine (Applied Biosystems) using the PowerUp SYBR green master mix (Applied Biosystems, Cat #A25742). Where possible, qPCR primers were designed to flank exon-intron junctions to prevent gDNA amplification. A table of primer sequences is included (Resource Table). Relative expression was calculated using the ΔΔCT method (Livak and Schmittgen, 2001). An average of *Hprt*, *Actb* and *Rplp0* expression levels were used for internal sample normalization.

All qRT-PCR experiments were carried out on three independently derived and treated mTEC populations. Significance testing was performed on log2 fold-changes in Prism 9, using an ordinary two-way ANOVA test with Holm-Sidak’s multiple hypothesis correction.

### Embryonic trachea collection

Trachea from embryos at day 17.5 of development (E17.5) were dissected into 4% paraformaldehyde in PBS. Trachea were incubated at 4°C for 2 hours. After fixative was washed off, samples were cryo-protected in 30% sucrose in PBS overnight at 4°C. Trachea were then embedded in OCT and stored at −80°C until 10-12 μm frozen sections were made with a Leica CM1900 cryostat. Sections were processed for RNA-FISH as described for mTECs above.

### RNA-FISH

Fluorescent in situ hybridization was performed using RNAscope Multiplex Fluorensence Reagent kit v2 (Advanced Cell Diagnostics, Cat #323100). mTEC cultures were washed three times in PBS and fixed for 15 min in 4% PFA in PBS. Fixed cells were washed three times with PBS for 5 min each and stored in PBS at 4°C for less than 1 week. mTEC transwell membranes were cut into six pieces and each piece was placed on a slide inside a hydrophobic boundary. RNAscope washes were performed on the slide within the hydrophobic boundary. All steps were performed in the HybEZ Humidity Control Tray. Both mTECs and embryonic trachea sections were washed twice with PBS and then treated with 30% hydrogen peroxide for 10 min at room temperature. After two washes in PBS, mTECs were incubated for 10 min at room temperature in Protease III diluted 1:15 in PBS whereas embryonic trachea sections were incubated for 15 min at 40°C with Protease III. Following two washes with PBS, RNAscope probes were added to mTECs and embryonic trachea sections for 2 hours at 40°C, washed twice with 1X RNAscope Wash buffer, and stored overnight in 5X SSC at room temperature. Amplification and HRP development steps were performed according to manufacturer’s instructions. Opal dyes were used at a dilution of 1:750 (Perkin Elmer, Cat #FP1488001KT and #FP1495001KT). To multiplex with immunostaining, RNA-FISH stained mTECs and embryonic trachea were blocked for 1 hour at room temperature in PBDT (1% BSA, 1% DMSO and 0.5% TritonX-100 in PBS) supplemented with 5% normal donkey serum and then incubated in primary antibody diluted in PBDT overnight at 4°C. Cells were washed three times in PBS for 5 min each and then incubated 1 hour at room temperature in secondary antibody diluted in PBDT along with Hoechst to stain nuclei. Cells were mounted in Prolong Diamond Antifade (Molecular Probes, Cat #P36970).

Fluorescent RNA *in situ* hybridization images were acquired on a Leica SP8 with spectral imaging capabilities. Three random fields of view per image was quantified manually for RNAscope probes. A minimum of three biological replicates were counted. Cells with 10 or more *Mycl* or *Pigr* RNAscope dots were considered positive for the respective marker. Categories of cells that represented less than 2% of cells were not included in graphs. Significance testing was performed in Prism 9 using multiple paired t-tests.

### CUT&RUN

CUT&RUN was performed as described (Meers et al., 2019; Peter J. Skene and Henikoff, 2017), with protocol and buffers detailed in Protocol 2: High Ca2+/low salt (https://www.protocols.io/view/cut-amp-run-targeted-in-situ-genome-wide-profiling-14egnr4ql5dy/v3). Briefly, MYCL-GFP, HEY1-GFP or NLS-GFP transduced mTECs were dissociated in 0.05% trypsin-EDTA in cell dissociation buffer for 15 min at 37°C. Cells were collected in in mTEC Basic medium with 2% Nu-Serum, washed twice in 5 mL of CUT&RUN Wash Buffer, and resuspended in CUT&RUN Wash Buffer. One million cells of each sample were aliquoted in 1.5 mL Eppendorf tubes and brought up to a volume of 1 mL with CUT&RUN Wash Buffer. Cells were incubated with 10 μL activated concavalin A beads (Bangs Laboratories, Cat #BP531) at room temperature for 10 min with gentle rocking. Cells were incubated overnight with anti-GFP (Abcam, Cat #ab290) diluted 1:100 in 0.05% digitonin in CUT&RUN Wash Buffer. Following two washes in CUT&RUN Wash Buffer, samples were incubated for 1 hour at room temperature in 1:100 rabbit anti-guinea pig IgG in 0.05% digitonin in CUT&RUN Wash Buffer. Samples were incubated with Protein A/G-MNase fusion protein (Addgene, Cat #123461) at 700 ng/mL for 1 hour at 4°C. Following two washes in 0.05% digitonin in CUT&RUN Wash Buffer and one wash in CUT&RUN Low-Salt Rinse Buffer, samples were digested for 10 min at 0°C in cold CUT&RUN Incubation Buffer. Chromatin fragments were released by incubation at 37°C for 30 min and then collected by phenol-chloroform extraction. Library preparation was performed with the NEB Next Ultra II DNA Library Prep Kit for Illumina (NEB, Cat #E7645S) using 25 μL of CUT&RUN DNA as input, as detailed by Liu N. (https://www.protocols.io/view/library-prep-for-cut-amp-run-with-nebnext-ultra-ii-kxygxm7pkl8j/v2). NEB adaptors were diluted 1:15 in adaptor dilution buffer. High sensitivity D5000 ScreenTape reagents (Agilent, Cat #5067-5593) and the Tapestation 4200 (Agilent) were used to assess library quality and pool libraries for sequencing. Sequencing was done on the Illumina NextSeq 500.

CUT&RUN analysis was performed based on previously described scripts (Zhu et al., 2019) and detailed in https://www.protocols.io/view/cut-amp-tag-data-processing-and-analysis-tutorial-e6nvw93x7gmk/v1. Briefly, FASTQs were trimmed using trimmomatic (Bolger et al., 2014) with a second trimming step to remove any remaining read-through adaptors. Reads were aligned with bowtie2 (Langmead and Salzberg, 2012) using parameters --local --very-sensitive-local --no-unal --no-mixed --no-discordant --phred33 -I 10 -X 700. Samples were filtered for reads under 120bp and duplicates were marked, but not removed, with picard MarkDuplicates. MACS2 (Zhang et al., 2008) was used to call peaks on merged replicates using parameters --keep-dup all -q 0.0001. The NLS-GFP controls samples from both MYCL and HEY1 experiments were merged to create a joint control sample and used as the control sample in the MACS2 callpeaks command to increase specificity of peak calling. ChIPseeker (Yu et al., 2015) was used to annotate peaks to the nearest gene. Peaks designated as ‘Promoter’ are within 3kb of the transcriptional start site (TSS). Peaks downstream of a gene, within the 5’UTR or 3’UTR or within a gene exon were combined into “Other”.

## Supporting information

Resource Table

## Statistical Testing

All statistical testing was performed using Prism 9 software. Statistical tests and multiple hypothesis corrections used for each experiment are listed under the relevant heading.

## Code Availability

Scripts are available at https://github.com/lb15/basal-to-multiciliated.

## Data Availability

scRNA-seq and CUT&RUN raw FASTQ files and processed data will be available on GEO. scRNA-seq data will be deposited on CellxGene.

## Author Contributions and Notes

L.E.B, J.F.R and S.P.C designed research, L.E.B, R.D, and S.P.C performed research, L.E.B performed bioinformatic analysis, L.E.B and S.P.C analyzed data; and L.E.B, J.F.R and S.P.C wrote the paper. J.F.R and S.P.C supervised the research.

J.F.R is a co-founder of Sen therapeutics and CorrectorBio.

This article contains supporting information online.

## Acknowledgments

We thank D. Erle, K.D. Koh and L. Bonser for help developing the CRISPR knockout protocol, E. Yu for assistance with mouse husbandry and members of the Reiter laboratory for critical discussion. We also thank N. Neff, R. Sit and M. Tan from the CZ Biohub Genomics platform for sequencing, the UCSF Laboratory for Cell Analysis for flow-cytometry machine use and assistance, the UCSF Center for Advanced Cell Technology for use of equipment and the Wynton high performance computing cluster for analysis. L.E.B was supported by Ruth L. Kirschstein National Research Service Awards (5T32HL007731-27 and 1F32HL154611-01). This work was supported by a CIRM Discovery grant (DISC1-10475) to S.P.C. and grants from the NIH (R01AR054396 and R01HD089918) to J.F.R.

