## Supplemental Figures 1-6 for "Opposing transcription factors MYCL and HEY1 mediate the Notch-dependent airway stem cell fate decision"

### Supplementary Figures

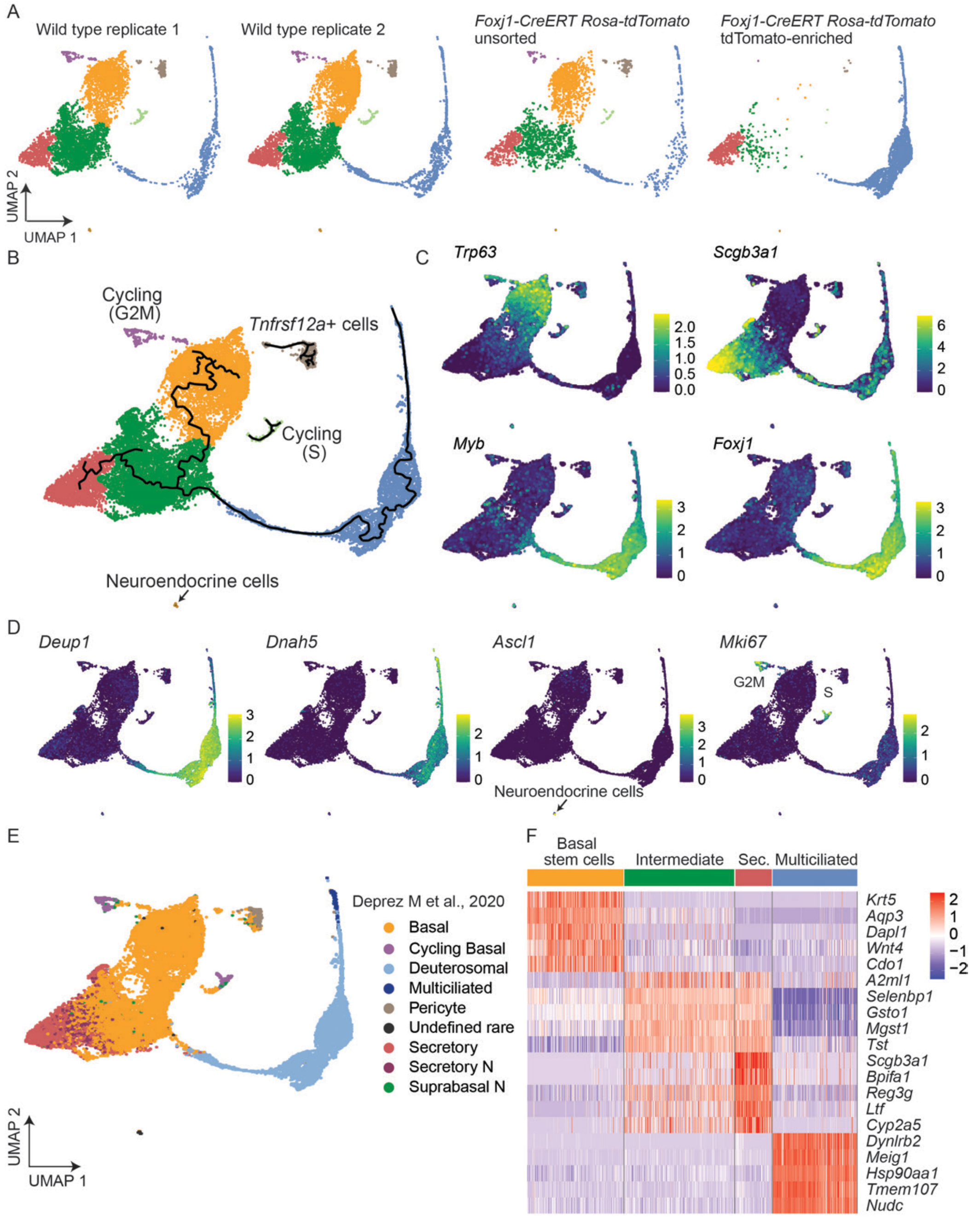


**Figure S1. scRNA-seq characterization of differentiating primary tracheal epithelial cells, related to Figure 1**

**A)** Contribution of each dataset to UMAP structure. Cells are colored by cluster identity. **B)** UMAP of scRNA-seq data generated from primary mouse tracheal epithelial cells three days after differentiation. Cells are colored by cluster identity and less common cell populations are labeled. Pseudotime paths are denoted as black lines. **C)** Expression of a basal stem cell marker gene (*Trp63*), a secretory cell marker gene (*Scgb3a1*) and multiciliated cell marker genes (*Myb* and *Foxj1*) overlaid on the UMAP. **D)** Expression of a marker of deuterosomal multiciliated lineage cells (*Deup1*), mature multiciliated cells (*Dnah5*), neuroendocrine cells (*Ascl1*) and cycling cells (*Mki67*) superimposed on the UMAP. **E)** Predicted cluster identities of mTECs differentiated for three days at air/liquid interface based on a human airway atlas. Cells are colored by cluster identity. **F)** Heat map of expression of the five most differentially expressed genes in basal stem cells, intermediate cells, multiciliated lineage cells, and secretory cells.

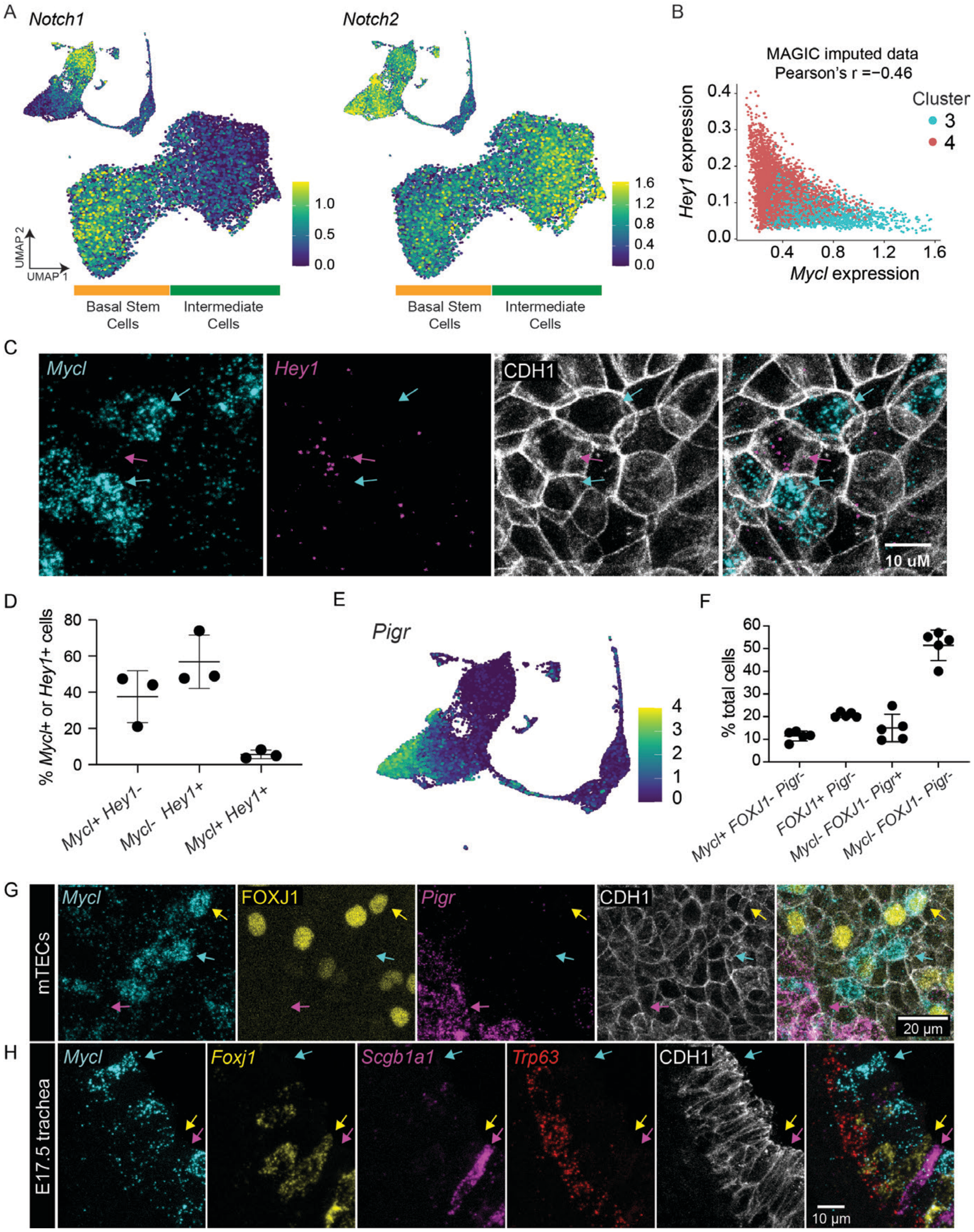


**Figure S2. Airway intermediate cells express *Mycl*, related to Figure 1**

**A)** Expression of Notch receptor genes *Notch1* and *Notch2* overlaid on UMAP of full dataset and UMAP of subclustered basal and intermediate cells. **B)** Pearson's correlation of *Mycl* and *Hey1* expression in clusters 3 and 4 of subclustered basal and intermediate cells. Correlation was performed on MAGIC imputed data. **C)** Visualization of *Mycl* and *Hey1* transcripts in mTECs differentiated for 3 days at air/liquid interface. Different subsets of intermediate cells expressed either *Mycl* (cyan arrows) or *Hey1* (magenta arrow). **D)** The proportions of *Mycl* and *Hey1*-expressing cells. Data are quantified from images in (C). Error bars represent standard deviation of the mean. n=3 replicates of independently derived mTECs. **E)** Expression of a secretory cell marker gene (*Pigr*) overlaid on the UMAP. **F)** The proportions of cells expressing *Mycl*, *Pigr* and FOXJ1. Data are quantified from multiplexed *in situ* hybridization and immunofluorescence imaging of mTECs differentiated for 3 days at air/liquid interface. Error bars represent standard deviation of the mean. n=5 replicates of independently derived mTECs. **G)** Visualization of *Mycl* transcripts with markers of multiciliated lineage cells (FOXJ1) and secretory cells (*Pigr*) in mTECs differentiated for 3 days at air/liquid interface. Cells expressed *Mycl* only (cyan arrow), both *Mycl* and FOXJ1 (yellow arrow), or *Pigr* only (magenta arrow). **H)** Multiplexed fluorescent *in situ* hybridization combined with immunofluorescence imaging of E17.5 trachea showing the expression of *Mycl* with a multiciliated cell marker (*Foxj1*), a secretory cell marker (*Scgb1a1*) and a basal stem cell marker (*Trp63*). E-cadherin (CDH1) immunofluoresence marked cell borders. Cells expressed *Mycl* only (cyan arrow), both *Mycl* and *Foxj1* (yellow arrow) or *Scgb1a1* only (magenta arrow).

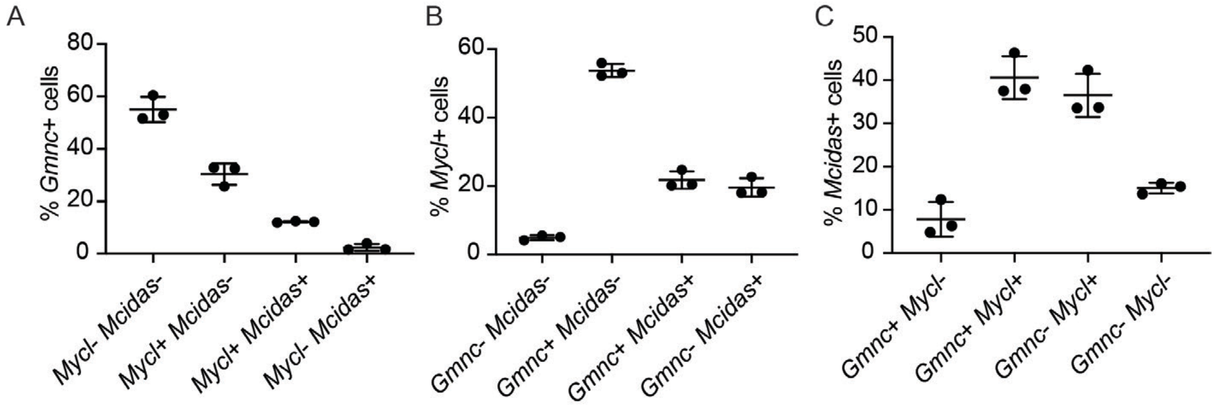


**Figure S3. *Mycl* is co-expressed with genes encoding regulators of early multiciliated cell differentiation, *Gmnc* and *Mcidas*, related to Figure 2**

**A)** The proportions of *Gmnc*-expressing cells also expressing *Mycl* or *Mcidas.* Data are quantified from images in Figure 2B. Error bars represent standard deviation of the mean. n=3 replicates of independently derived mTECs. **B)** The proportions of *Mycl*-expressing cells which also expressing *Gmnc* or *Mcidas.* Data are quantified from images in Figure 2B. Error bars represent standard deviation of the mean. n=3 replicates of independently derived mTECs. **C)** The proportions of *Mcidas*-expressing cells also expressing *Gmnc* or *Mycl.* Data are quantified from images in Figure 2B. Error bars represent standard deviation of the mean. n=3 replicates of independently derived mTECs.

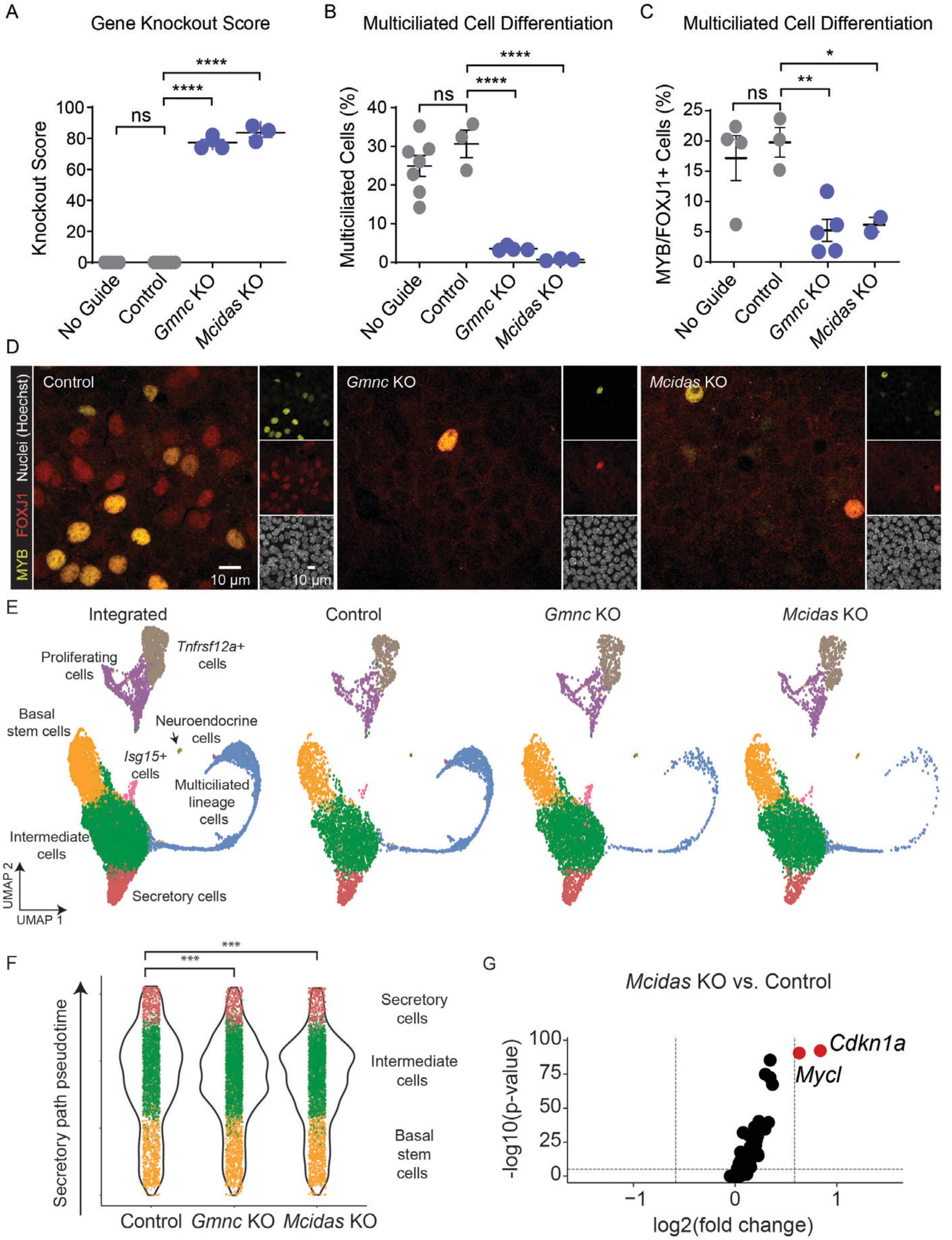


**Figure S4. *Gmnc* and *Mcidas* are dispensable for the initiation of the multiciliated lineage, related to Figure 3**

**A)** Quantification of knockout efficiency of *Gmnc* or *Mcidas* in mTECs receiving no gRNA, a non-targeting gRNA (control), a gRNA targeting *Gmnc* or a gRNA targeting *Mcidas* after five days at air/liquid interface. Knockout score indicates the percentage of frame-shift mutations predicted from Sanger sequencing PCR-amplified region around the targeted locus (see Methods for further details). Significance was assessed using a one-way ANOVA test with multiple comparison correction, with ****

indicating p < 0.0001. Error bars represent SEM for replicates of independently derived and electroporated mTECs. **B)** Quantification of the percentage of multiciliated cells among mTECs receiving no guide RNA, a non-targeting guide RNA (control), or a guide RNA targeting *Gmnc* or *Mcidas* after 21 days at air/liquid interface. Significance was assessed using a one-way ANOVA test with multiple comparison correction, with **** indicating p < 0.0001. Error bars represent SEM for replicates of independently derived and electroporated mTECs. **C)** Quantification of the percentage of MYB+/FOXJ1+ cells among mTECs receiving no guide RNA, a non-targeting guide RNA (control), or a guide RNA targeting *Gmnc* or *Mcidas* after five days at air/liquid interface. Significance was assessed using a one-way ANOVA test with multiple comparison correction, with ** indicating p = 0.0056, while * indicates p < 0.0308. Error bars represent SEM for replicates of independently derived and electroporated mTECs. **D)** Immunofluorescence images of mTECs receiving no guide RNA, a non-targeting guide RNA (control), or a guide RNA targeting *Gmnc* or *Mcidas*. After five days at air/liquid interface, cells were immunostained for MYB and FOXJ1. **E)** UMAP of integrated control, *Gmnc* KO, and *Mcidas* KO scRNA-seq data. Clusters are colored by cell identity (left). Contribution of each dataset to UMAP structure (three right panels). **F)** Violin plot of pseudotime values along the secretory differentiation path for each cell. Cells are colored by clusters as depicted in (Fig. 3B). ***Differential progressionTest (condiments R package) p-value < 0.005. **G)** Volcano plot of *Mycl*-correlated genes in *Mcidas* KO vs. control intermediate cells. Dashed line demarcates 1.5-fold increase in expression. *Mycl* and *Cdkn1a* are expressed more than 1.5-fold higher in *Mcidas* KO intermediate cells (red).

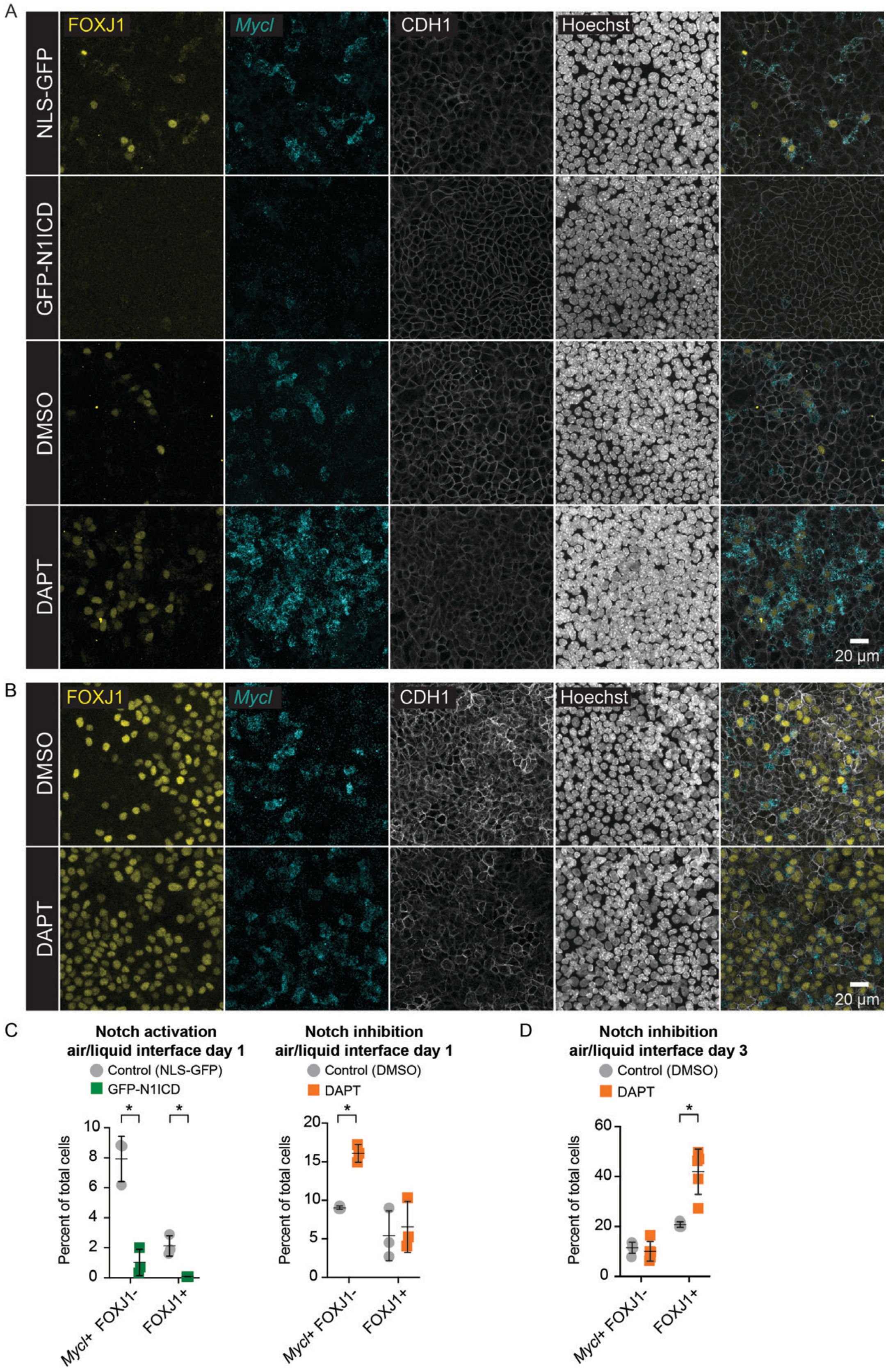


**Figure S5. Notch signaling represses *Mycl* expression, related to Figure 5**

**A)** Multiplexed RNA-FISH and immunofluorescence of mTECs cultured for one day at air/liquid interface with inhibition of Notch signaling (treated with 1μM DAPT) or activation of Notch signaling (transduced with GFP-N1ICD). Controls are vehicle treated (DMSO) or NLS-GFP transduced, respectively. *Mycl* staining was multiplexed with Hoechst and immunostaining for FOXJ1 and CDH1. **B)** Multiplexed RNA-FISH and immunofluorescence of mTECs cultured for three days at air/liquid interface with inhibition of Notch signaling (treated with 1μM DAPT) or control (treated with DMSO). *Mycl* staining was multiplexed with Hoechst and immunostaining for FOXJ1 and CDH1. **C)** The proportions of cells expressing *Mycl* and FOXJ1 upon Notch activation or abrogation in mTECs cultured for one day at air/liquid interface. Data are quantified from images in (A). Significance was assessed using multiple paired t tests with multiple comparison correction, with * indicating $p < 0.05$. Error bars represent standard deviation of the mean for 3 replicates of independently derived and transduced mTECs. **D)** The proportions of cells expressing *Mycl* and FOXJ1 upon Notch activation in mTECs cultured for three days at air/liquid interface. Data are quantified from images in (B). Significance was assessed using multiple paired t tests with multiple comparison correction, with * indicating $p < 0.05$. Error bars represent standard deviation of the mean for 5 replicates of independently derived and transduced mTECs.

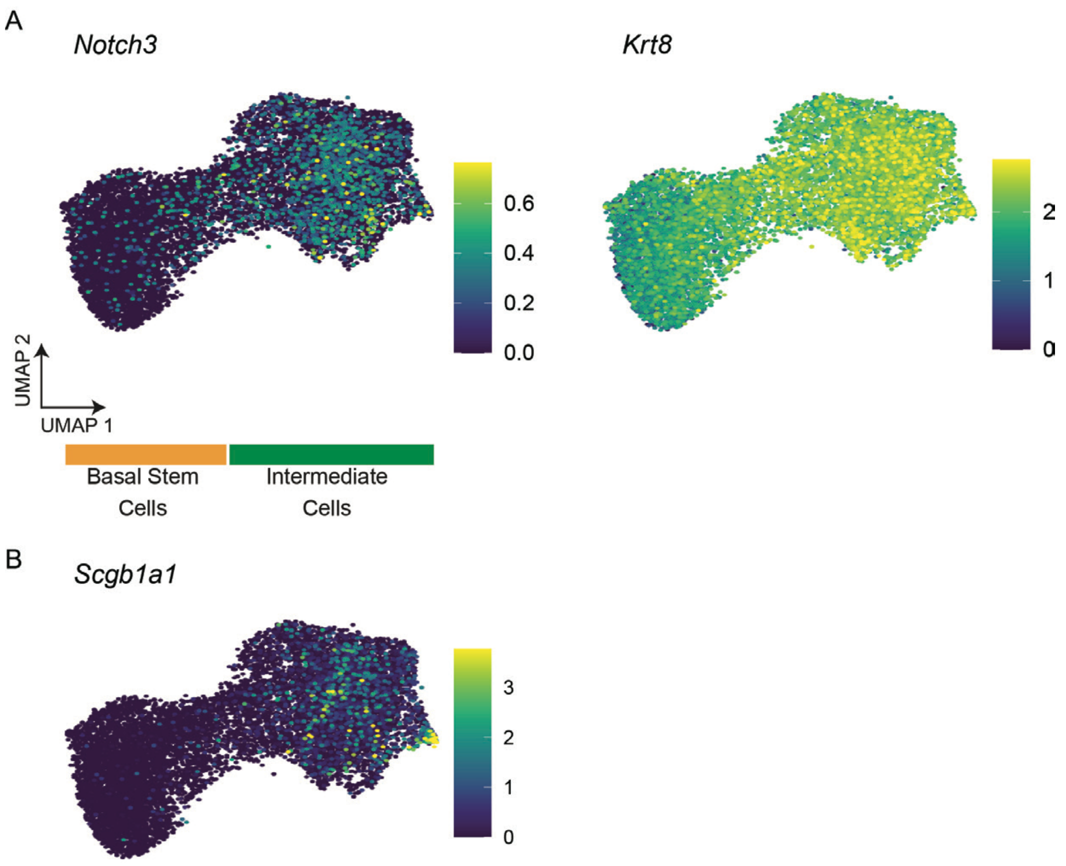


**Figure S6. Suprabasal and club cell markers are expressed in intermediate cells, related to Figure 7**

**A)** Expression of *Notch3* and *Krt8* in basal and intermediate clusters. **B)** Expression of *Scgb1a1* in basal and intermediate clusters.
